# Multi-task Intracranial EEG recordings reveal a Comprehensive Role of the Human Dorsal Anterior Insula in High-Level Cognition

**DOI:** 10.64898/2025.12.15.694298

**Authors:** Benoit Chatard, Maryne Dupin, Mathilde Petton, Benjamin Bontemps, Lorella Minotti, Philippe Kahane, Sylvain Rheims, Julien Bastin, Jean-Philippe Lachaux

## Abstract

The dorsal anterior insula (dAI) is widely recognized as a cornerstone of human cognition. However, its functional characterization remains the subject of considerable debate and competing theories. Embracing a “shift in viewpoint” approach—defining a brain region’s function through the observation of the same neural population across diverse experimental contexts—this study used intracranial EEG (iEEG) recordings from a large cohort of patients to investigate the sub-second neural response dynamics of the human dAI across a broad range of cognitive tasks, spanning visual category discrimination, phonological and semantic processing, attentive visual search, and verbal and visuospatial working memory. This within-subject, multi-task iEEG framework offered a robust basis for decoding the structure–function relationship of the dAI and for empirically testing key predictions of prevailing theoretical models concerning its role. Our findings confirmed the dAI’s involvement in the Action-Mode Network and shed new light on the large-scale spatiotemporal organization of that network, while providing novel insights into the interplay between cognitive effort and efficiency.

## Introduction

The insula lies parallel to the interhemispheric plane within the fundus of the Sylvian sulcus, beneath the temporal and frontal lobes. Its characteristic seashell-like morphology is divided by the central insular sulcus into posterior and anterior subregions that markedly differ in terms of cytoarchitectonic organization, as well as structural and functional connectivity (***Augustine (1996); Evrard (2019); Flynn (1999); Namkung et al. (2017); Quabs et al. (2022))***. While the human posterior insula is predominantly involved in interoception – monitoring the body’s internal state – and processing proximal environmental stimuli, including chemosensation, as in many species (***Craig (2009); Craig (2010); Mazzola et al. (2019); Namboodiri et al. (2015); Segerdahl et al. (2015); Small (2010))***, the human anterior insula (AI) is characterized by a disproportionate evolutionary expansion to support cognitive processes related to the abstract integration of past and present states to assess situations and guide forthcoming behavior, including social behavior (***Bastin et al. (2017); Billeke et al. (2020); Centanni et al. (2021)***; ***Craig (2009); Damasio (2003); Droutman et al. (2015); Gogolla (2017)***; ***Molnar-Szakacs and Uddin (2022)***; ***Nelson et al. (2010))***. The AI has been considered for decades as an enigmatic “multi-purpose swiss-knife” of higher cognition, because of its systematic activation in a broad spectrum of neuroimaging paradigms (***Kurth et al., 2010***). But since 2010, a finer understanding has emerged emphasizing a clear functional dissociation between the ventral anterior insula (vAI), preferentially involved in socio-emotional processes, and the dorsal anterior insula (dAI) associated with high-level cognitive functions such as attention, task-level performance monitoring, decision-making or perceptual awareness (***Centanni et al. (2021)***; ***Llorens et al. (2023)***; ***Molnar-Szakacs and Uddin (2022))***.

Yet, many uncertainties persist, especially regarding the dAI, that several opposing theories currently associate with widely distinct functions ranging from perceptual awareness or error prediction, to saliency detection or task-level cognitive control (***Billeke et al. (2020***); ***Citherlet et al. (2020***); ***Han et al. (2018***); ***Huang et al. (2021***); ***Llorens et al. (2023***); ***Nelson et al. (2010***)). And while there is a broad agreement on its pivotal role in processing behaviorally-relevant stimuli, the specific stage of its contribution remains a subject of intense debate. While some authors have proposed that the dAI essentially selects stimuli for further processing (pure saliency detection), others have suggested a more active role in the initiation and the orchestration of the cognitive response to selected stimuli (***Dosenbach et al. (2008***); ***Dosenbach et al. (2025***); ***Menon (2011***); ***Menon et al. (2020***); ***Molnar-Szakacs and Uddin (2022***); ***Uddin (2015***)). Part of this ongoing confusion is likely attributable to the inherent limitations of neuroimaging data upon which most current theories are based, as fMRI notoriously blurs the distinction between spatially and temporally proximal cognitive processes (***Menon, 2011***).

Direct neural recordings of the human dAI, using intracerebral electroencephalography (iEEG) in neurological patients, offer a unique opportunity to bypass the technical limitations of fMRI (***Lachaux et al. (2012***); (***Mercier and others (2022***)), and several iEEG studies of the insula have already yielded valuable insights into its regional response dynamics during single cognitive tasks with great anatomical and temporal specificity (***Bastin et al. (2017***); ***Billeke et al. (2020***); ***Blenkmann et al. (2019***); ***Cecchi et al. (2022***); ***Citherlet et al. (2019***); ***Das and Menon (2020***); ***Das and Menon (2024***); ***Duong et al. (2023***); ***Gueguen et al. (2021***); ***Llorens et al. (2023***)). However, a comprehensive functional characterization of a brain region like the dAI requires data recorded in a variety of cognitive situations, ideally within the same neural populations in the same brains (***Genon et al., 2018***)). The present study shows that iEEG recordings collected across multiple cognitive tasks with high functional, anatomical and temporal precision in the same subjects, help refine our understanding of the dAI.

We recorded intracranial EEG in 169 participants while they performed a set of cognitive tasks including visual category discrimination, phonological and semantic processing, attentive visual search and verbal and visuo-spatial working memory. Our analysis focused on the detailed spatial-temporal organization of the dAI response across tasks to test implicit predictions of existing theories regarding the temporal processing stages of that region. Altogether, our findings provide novel insights into the fundamental role of that cornerstone of human cognition, revealing a more comprehensive contribution to high-level cognition than most recent theories have proposed.

## Results

### A sustained neural response in the dorsal anterior insula during task-performance

Using the HiBoP software (***Vecchio et al., 2024***), a total of 621 intracranial EEG (iEEG) recording sites located within or in the immediate vicinity of the insula were identified and labeled based on the individual anatomy of 140 patients (and 16553 sites) (***Figure 1***). A preliminary examination of the High-Frequency Activity (HFA) time-courses in those sites during our experimental tasks (***Figure 1***) (visible in trial matrices, ***Figure 2***), revealed a pattern consistent with the dichotomy reported by Llorens et al. (***Llorens et al., 2023***), with a sustained HFA increase during the stimulus-response interval, predominantly in the anterior insula, and a distinct pattern characterized by a post-response peak, primarily in the posterior insula. To formally dissociate those apparent clusters, hierarchical clustering was applied to the 621 sites, using a condensed quantification of the HFA time-courses relative to stimulus and response onsets (***Figure 2***) in the MCSE-HARD and LEC1-SEMA experimental conditions (which were chosen because they involved very different cognitive processes and a large number of behavioral responses, and because including more conditions would have exaggeratedly increased the dimensionality of the clustering space). The analysis confirmed the sharp dissociation between a) a cluster of 294 sites, mostly located in the anterior part of the insula (***Figure 3***) with a sustained neural activity during the stimulus-response interval, and b) two additional clusters (with 200 and 127 sites respectively) predominantly posterior, with an activity that was either weak or mainly followed the motor response (e.g. posterior insular site shown in ***Figure 2***)). Given our interest in insular regions involved in task-performance, we focused subsequent analyses on the first cluster, which homogeneous dynamics was compatible with an active participation in the cognitive processes supporting stimulus processing, decision-making and response preparation (this cluster will be referred to as the “Task-Responsive Cluster”, or TR-CLUS, hereafter). A projection onto the cytoarchitectonical atlas of (***Quabs et al., 2022***) (***Figure 3***), revealed that a majority of TR-CLUS sites were located in areas ld6 or ld7 of the insula. Within that cluster, the cluster-level HFA time-course relative to stimulus and response time (illustrated by grand average trial matrices computed across sites and participants fo each task) were remarkably similar across all experimental conditions of MCSE and LEC1, and characterized by a sustained HFA increase reaching its peak prior to the motor response, compatible with an active role in the perception-decision-action sequence. Observing such a similar pattern across largely dissimilar tasks suggested a task-general, rather than task-specific, function for that cluster.

**Figure 1.**
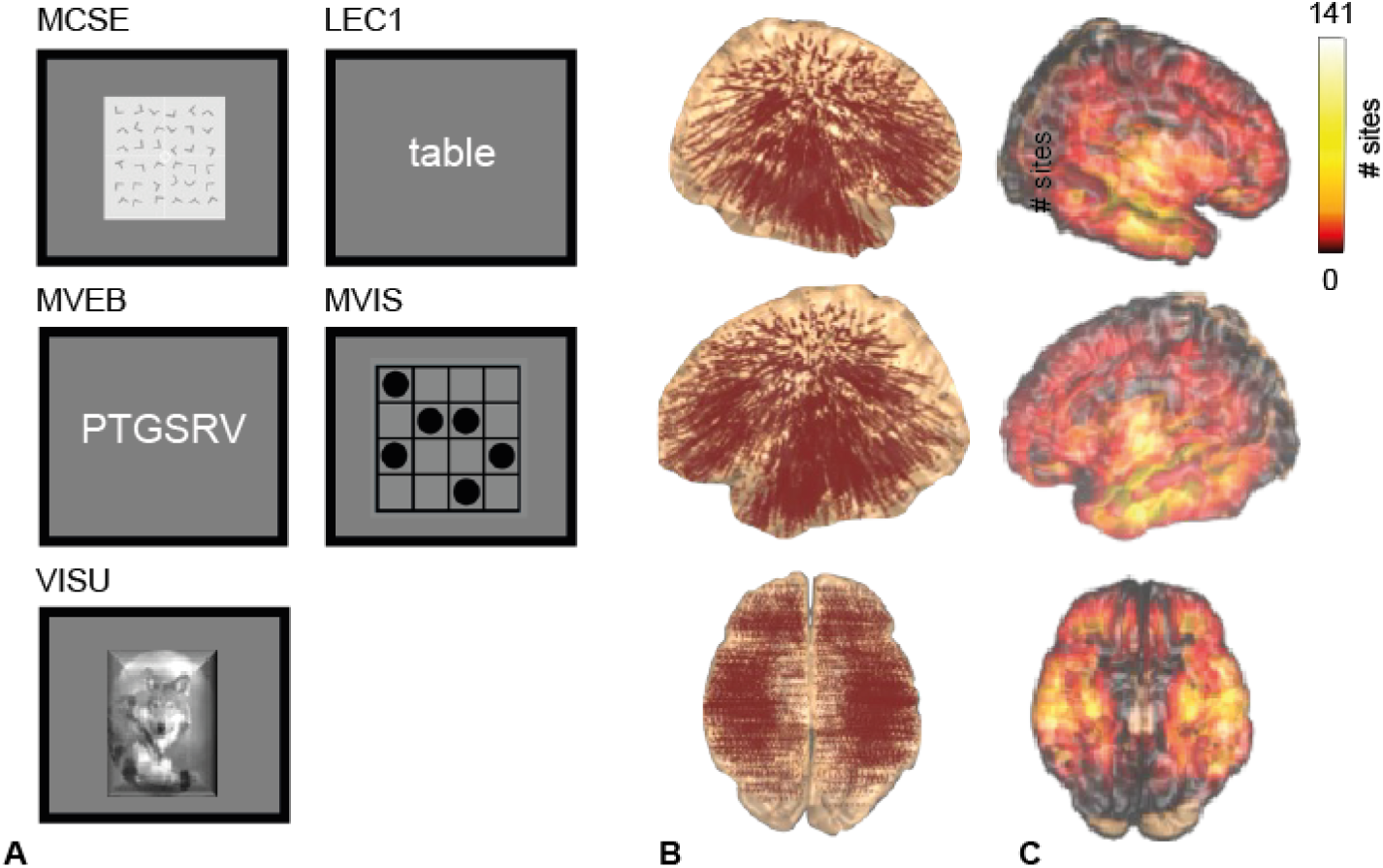
Overview of the experimental paradigms and iEEG brain sampling. (A) Representative stimuli used in the five experimental paradigms (MCSE, LEC1, MVEB, MVIS and VISU; see methods for detailed descriptions). (B) Spatial distribution of all recorded iEEG electrode sites across all participants, visualized on a standard MNI brain template. Each marker represents a single electrode site. (C) Anatomical coverage showing the cumulative number of recording sites per brain region based on a 3D MNI brain parcellation. Regions are color-coded by site count.

**Figure 2.**
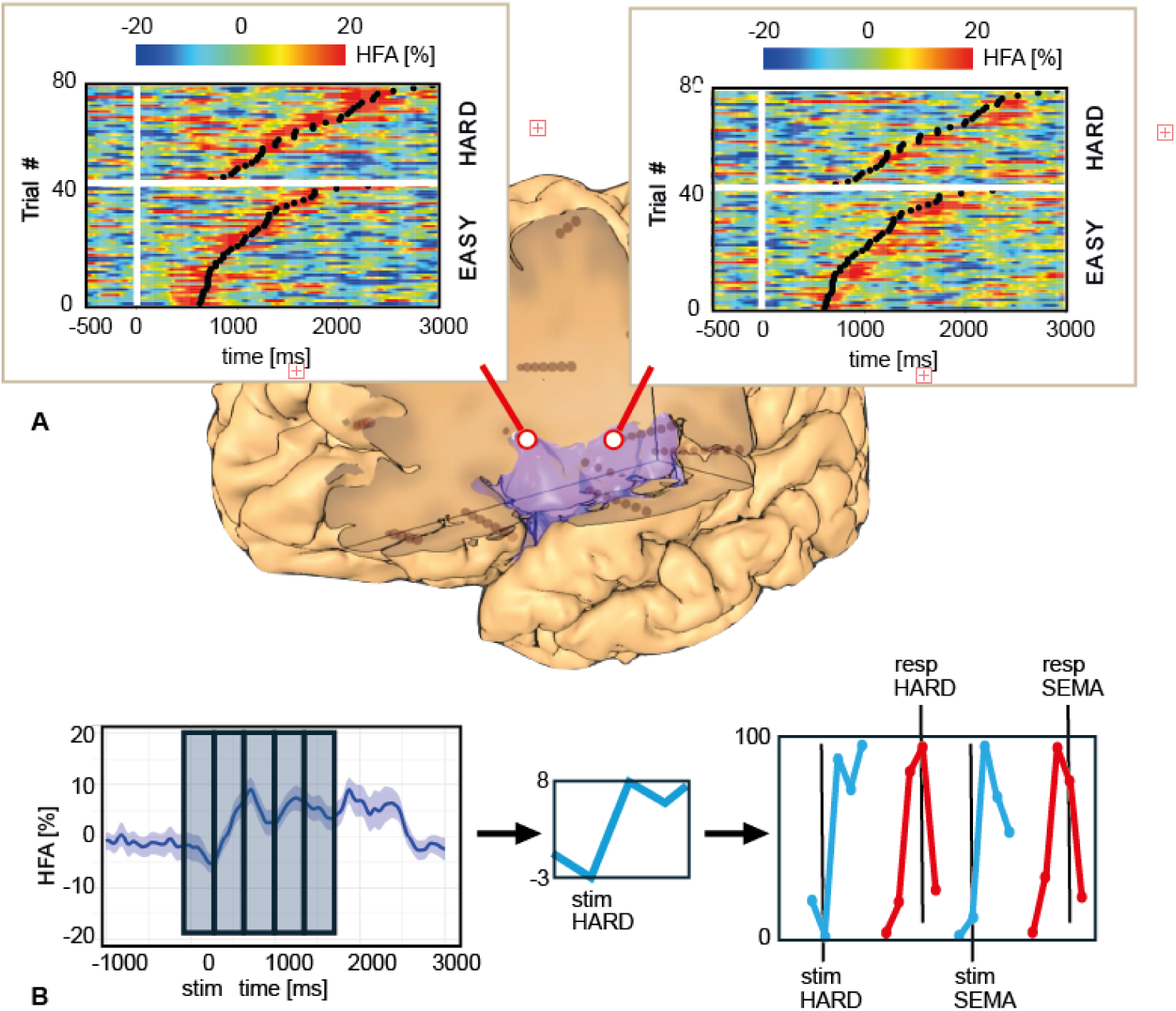
Temporal dynamics in the insula during visual search and data feature extraction for clustering. (A) Representative examples of the dynamics of High-Frequency Activity (HFA, 50-150 Hz) in the anterior and posterior insula of a single patient during the visual search task. Each matrix displays HFA over time (x axis) for all trials (y axis), sorted by reaction time (black oblique lines) for the EASY condition (lower section) and the HARD condition (upper section). Stimulus onset is at time 0 (white vertical line) and HFA is expressed in % change relative to the mean HFA across the entire experiment. Note that HFA peaks before (resp. after) reaction time in the anterior (resp. posterior) insula. (B) Data feature extraction method applied prior to hierarchical clustering of iEEG sites. HFA is epoched around a specific event (in this example, the stimulus onset of the MCSE HARD trials), then averaged across trials (left plot). The mean across five consecutive time-windows (vertical rectangles) provides five values (middle plot), which are normalized between 0 and 100. Epoching the MCSE HARD or the LEC1 SEMA trials relative to the stimulus onset or the response time provides a 20-dimensional vector for a given site (5 windows x 2 experimental conditions x 2 central epoching events).

**Figure 3.**
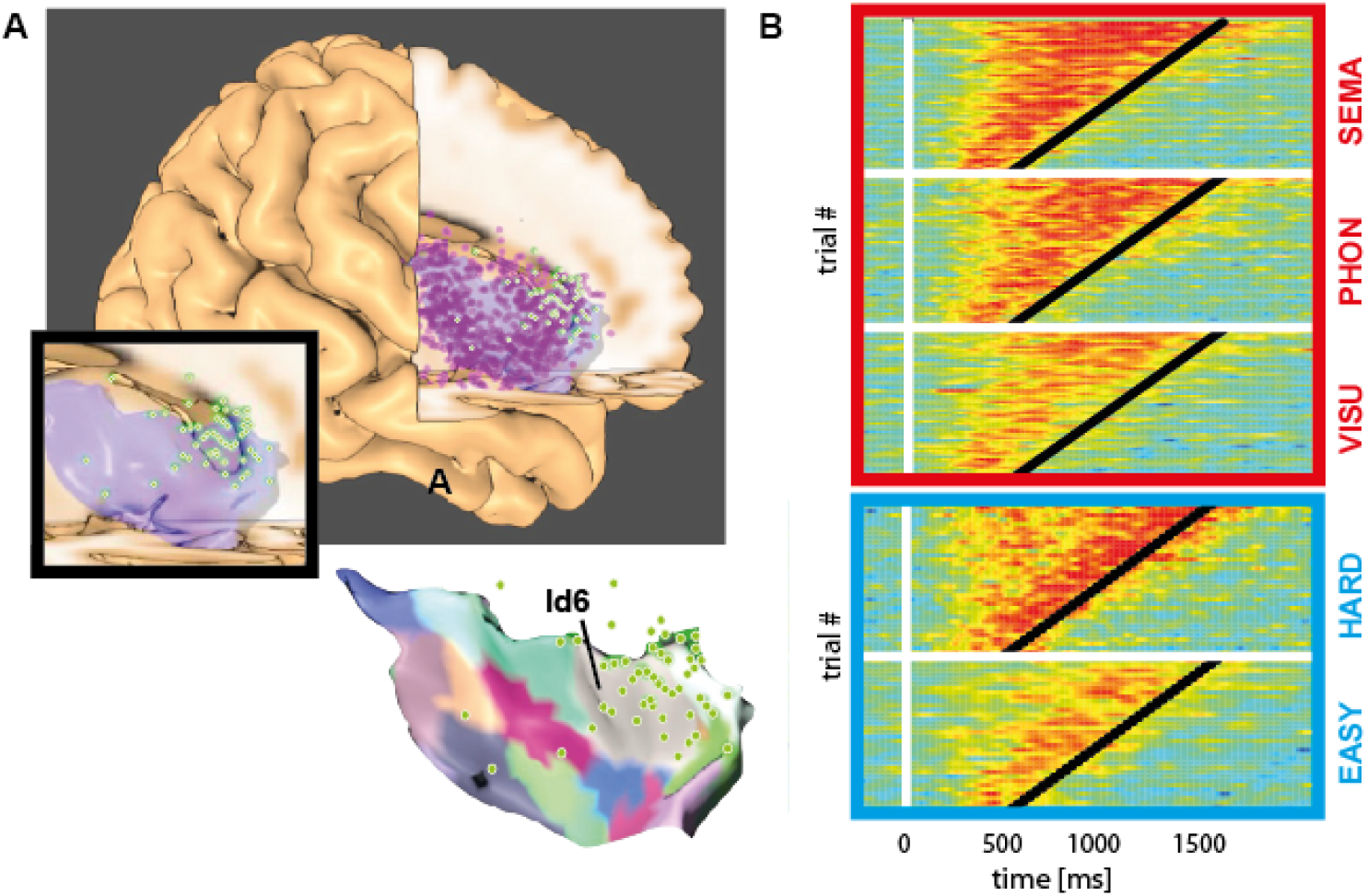
iEEG sampling of the insula and HFA dynamics of the dAI cluster. (A) Anatomical distribution of all iEEG sites in the insula, displayed on a 3D view of the MNI template (all sites have been projected onto the right insula). Sites in green correspond specifically to the first cluster, characterized by a strong HFA increase prior to reaction time (e.g. the anterior insular site in (***Figure 2***). The lower inset provides a magnified view of this dAI cluster (green sites, 294 sites in 80 patients), shown without (left) and with (right) the cytoarchitectonic parcellation by (***Quabs et al., 2022***), highlighting cytoarchitectonic area Id6 in grey. Note that the projection method can cause some insular sites to appear outside the template insula. (B) Average temporal dynamics of High-Frequency Activity (HFA, 50-150 Hz) across all sites within the dAI cluster (green sites) for all conditions of the MCSE and LEC1 tasks. Before averaging, differences in reaction time (RT) across participants were compensated using a time-stretching algorithm to fit a common 500-1500 ms RT distribution (see methods for details). The x-axis represents normalized time relative to stimulus onset.

### Anatomical specificity of the domain-general insular response

To reach more precise conclusions regarding that insular subdivision, we selected iEEG sites within TR-CLUS which matched the following anatomy-functional criteria: a) a confirmed localization within the insula (verified by visual inspections of each individual brain) and b) a significant neural response during both the MCSE-HARD and the LEC1-SEMA conditions (as defined by a significant HFA increase in a [500:1000 ms] post-stimulus window relative to the 200 ms pre-stimulus base-line, Wilcoxon test, p<0.05, FDR-corrected). This approach aimed to precisely delineate insular subregions actively engaged throughout those two separate, demanding cognitive situations, in the interval leading to the behavioral response. Out of 294 sites in TR-CLUS, 59 sites met both our anatomo-functional criteria. The majority of them (91%) were located in the anterior short gyrus (ASG, 34/59), the middle short gyrus (MSG, 13/59) and the posterior short gyrus (PSG, 7/59) even though the three short gyri comprised only 34% of the 621 iEEG sites included in the clustering analysis). Most notably, a vast majority of those 59 sites (80%) were located in the dorsal ASG (dASG) or dorsal MSG (dMSG, with a dorsal/ventral distinction based on the delineation between the upward/downward-facing insular surface), approximatively overlapping cytoarchitectonic ar-eas ld6 and ld7 (while representing only 27% of the iEEG across all three clusters). And while the number of iEEG sites recorded in the dASG (92) and the dMSG (76) were comparable, a substantially larger proportion of the 59 sites were found in the dASG (34 vs 13), suggesting a predomi-nant contribution of ld7 to task-performance. Conversely, we tried to evaluate the probability that a random site recorded in ld6 and ld7 would match our two anatomo-functional criteria. For that purpose, we considered all the depth-electrodes (i.e. SEEG linear arrays) traversing either the dASG or dMSG (60 depth-electrodes) and found that 56 included at least one iEEG site in TR-CLUS, 40 of which met our anatomy-functional criteria. Further examination of the remaining 16 electrodes revealed that a majority of them (13/16) included at least one TR-CLUS site with a significant response in either LEC1-SEMA or MCSE-HARD and that in most cases, an HFA peak was observed in the “non-significant” condition beyond our chosen time window (i.e. later than 1000 ms). Overall, our observations demonstrate a very strong functional homogeneity within the dAI (particularly within ld6/ld7).

### Dorsal-Anterior and Left-Right differentiation

In some patients, oblique depth-electrodes in the anterior insula provided some insight into the dorso-ventral extent of the task-responsive cluster. As illustrated by two representative cases in (Supplementary Figure 1), iEEG sites positioned ventrally, approximately one centimetre or more from the circular sulcus displayed a marked decrease of the observed task-response. This pattern of attenuation is consistent with the vertical differentiation suggested by microstructural parcellations, such as reported by Quabs and colleagues (***Quabs et al., 2022***). In addition, two patients with a bilateral implantation of the dAI allowed us to test for a possible inter-hemispheric specialization of the dAI (Supplementary Figure 2). Although it was impossible to draw significant conclusions from such a small sample, we found no evidence for a meaningful functional asymmetry of the dAI, at least in our tasks.

### Multi-functional signature of the dorsal anterior insula

To further characterize the functional signature of the dAI, and since consecutive sites along the same electrode often display similar response profiles, we selected the most responsive iEEG site on each of the 40 electrodes identified in the previous section (which correspond to 60 electrodes minus 4 with no TR-CLUS site and 16 lacking a significant response in either MCSE or LEC1). All of those 40 selected iEEG sites were located in the dAI, in TR-CLUS and had significant responses in MCSE-HARD and LEC1-SEMA (26 were located in the dASG, 8 in the dMSG and 4 in the dPSG, in 30 patients). That subset of dAI recordings provided important insights into the functional signature of this region across our five tasks (as defined by the combined task-related activation profiles measured in those tasks). ***Figure 4*** displays the average response profiles for that subset for all experimental conditions, together with similar response profiles observed in individual representative sites in ***Figure 5*** and Supplementary Figure 3. Specifically for the LEC1 and MCSE tasks, and in line with the trial-matrix depiction of ***Figure 3***, neural responses started within 500 ms after stimulus presentation and were longer in conditions that required a more sustained cognitive engagement (e.g. MCSE HARD vs. EASY, and LEC1 SEMA vs. CASE). The dAI also exhibited a robust activation during the working memory paradigms (MVEB and MVIS) independent of the type of stimuli to be encoded and stored (verbal or visuo-spatial). We confirmed previous findings (***Llorens et al., 2023***) that this region strongly reacts to the probe onset, but we also showed that dAI activity could increase endogenously, in the absence of external stimuli, during the initial phase of the maintenance period. That endogenous activity increased with memory load for both the verbal and visuospatial working memory paradigms: the comparison between the HFA measured in the initial stage [+1600 ms: 2400 ms] of the maintenance window (starting at 1500 ms) for the low load (2 items) and high-load conditions revealed a greater activity for the later in 29/40 sites for MVEB and 16/40 sites for MVIS (wilcoxon test, p<0.05, FDR correction). Results for the visual oddball task (VISU) confirmed the reported activation of the dAI in response to rare, task-relevant stimuli (***Citherlet et al., 2019***), as 39 out of 40 sites showed a significant HFA increase for the target stimuli (“fruits”, Wilcoxon comparison with pre-stimulus baseline, FDR-corrected p<0.05). But we also observed a weaker but significant response to several specific categories of non-target images, including animals (21/40 sites), objects (12/40 sites), landscapes (6/40) and pseudowords (5/40) (see ***Figure 5*** for a representative response to “animals”). Interestingly, face stimuli, which are very salient for the human species, elicited a response in only one dAI site, fewer than scrambled pictures (2/40).

**Figure 4.**
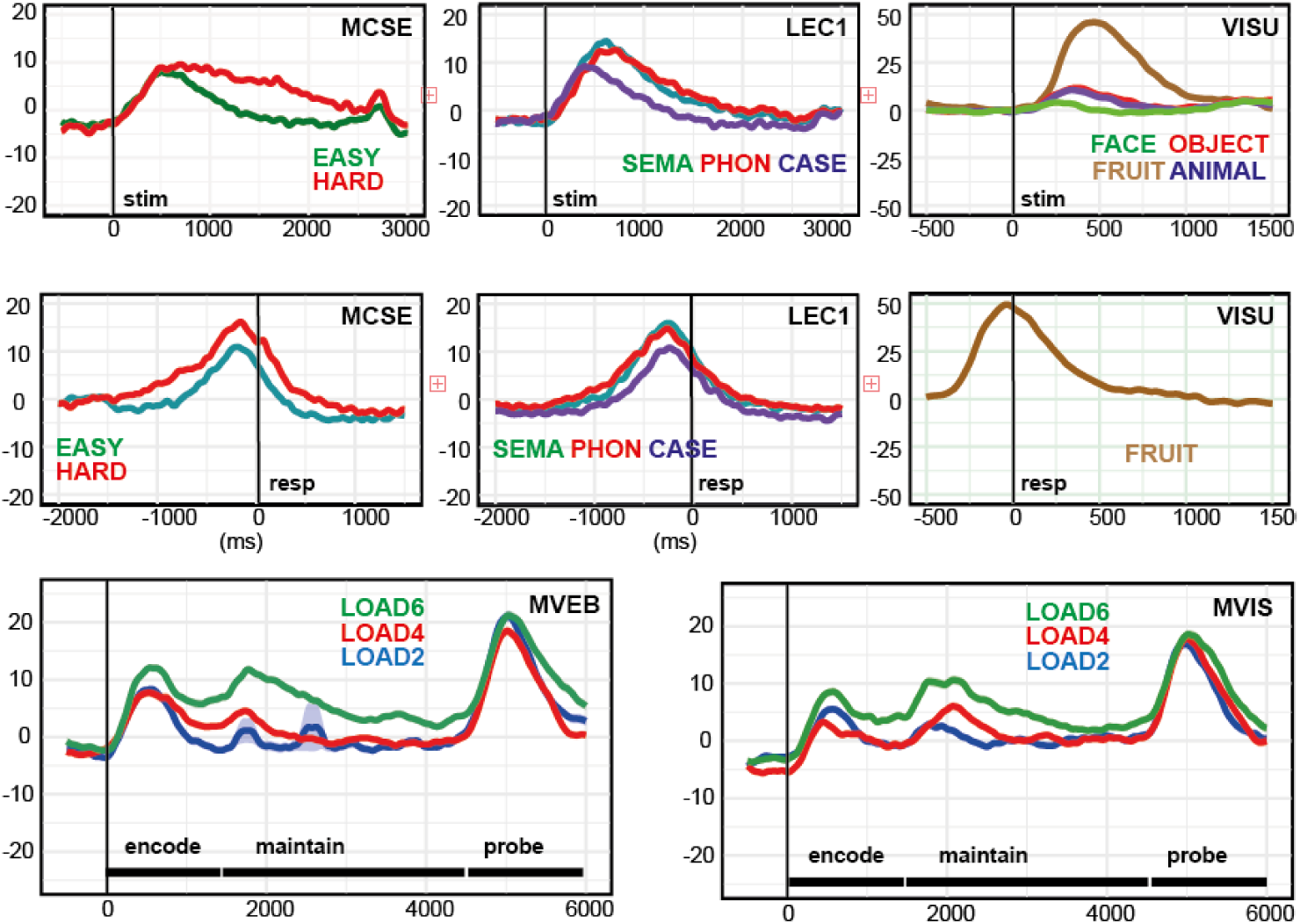
Multi-task characterization of the dorsal anterior insula functional response. Each graph depicts the High-Frequency Activity (HFA, 50-150 Hz), averaged across all dAI recording sites (green sites in ***Figure 3***, 294 sites in 80 patients) (expressed as percentage change relative to the mean HFA across the entire experiment, with standard error of the mean (s.e.m.)). Color codes experimental conditions. In the top row (resp. middle row) HFA was epoched in each trial relative to the stimulus onset, at 0 ms (resp. relative to the motor response). Note that in the VISU paradigm, only fruits stimuli were followed by a motor response. bottom row: average HFA response aligned to stimulus onset (0 ms) for the three levels of memory load in the two working memory protocols (MVEB and MVIS).

**Figure 5.**
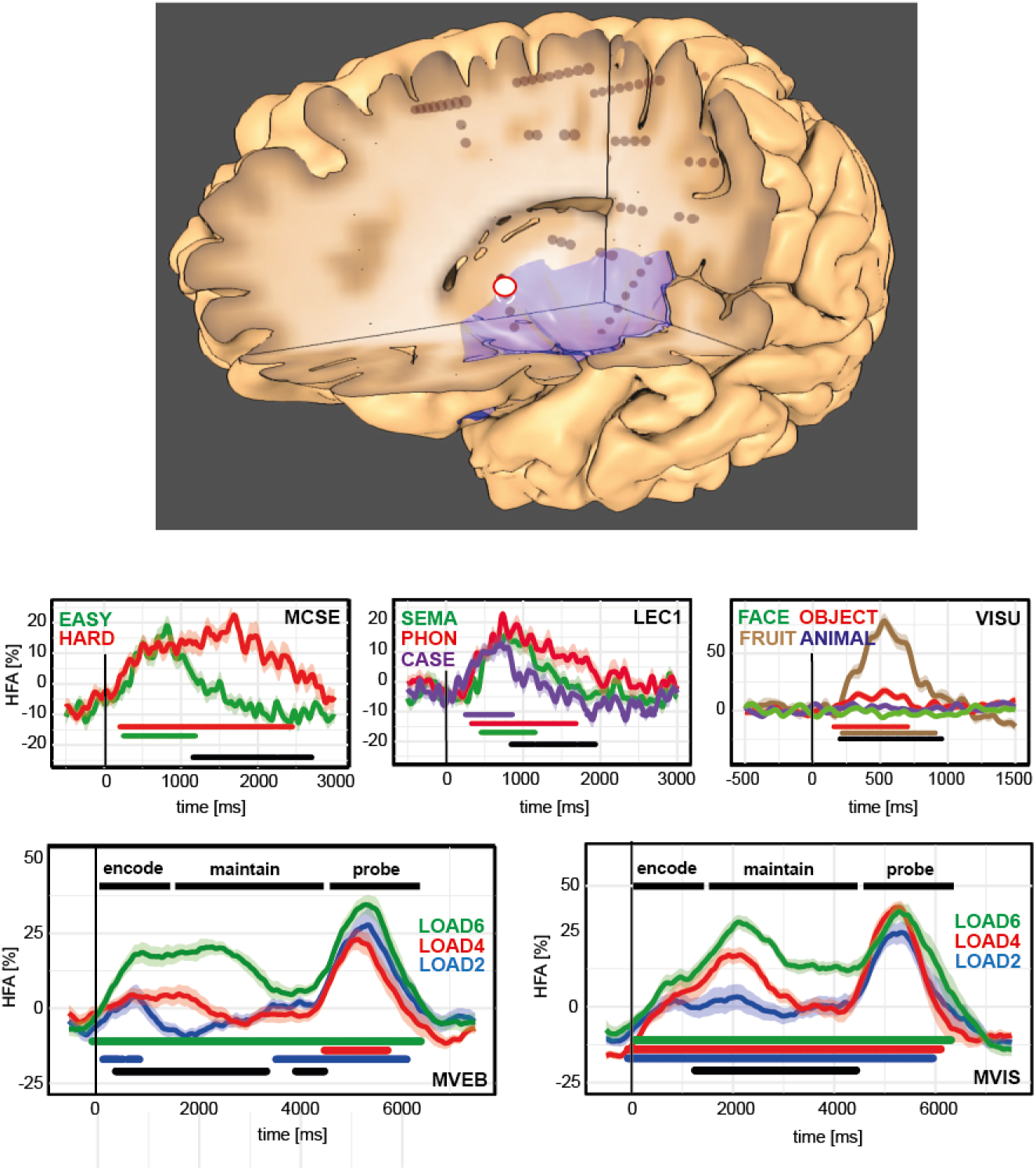
Representative example of the multi-task functional response in the dorsal anterior insula. Similar representation as in ***Figure 4***, for a single site in the superior portion of the short middle gyrus of the insula. Colored horizontal bars indicate time periods during which HFA is statistically higher than baseline (t-test, p<0.05, uncorrected, removing isolated intervals shorter than 100 ms). Black horizontal bars indicate time periods of statistically significant HFA difference between conditions (Kruskal-Wallis test, p<0.05, uncorrected). The white dot indicates the precise location of the recording site onto the participant’s individual anatomy.

### Later and stronger responses for longer reaction times

Building upon visual observations (***Figure 3***) suggesting that longer reaction times (RT) might be associated with a later HFA peak latency and potentially stronger HFA activity prior to the motor response, we formally tested these hypotheses across sites in the dAI 40-sites subset for the LEC1 and MCSE tasks, separately. We analyzed the trial-to-trial relationship between RT and either a) the HFA peak latency (“PeakLat”) or b) the mean HFA during the 400 ms window preceding the motor response (“PreRespMean”). A first analysis was performed on each site of our dAI subset individually while pooling the correct trials of all experimental conditions (i.e. HARD and EASY), to estimate the proportion of sites exhibiting a significant relationship between the HFA time-course and reaction time globally, during the entire task. We grouped correct trials into ten deciles based on RT from fastest to slowest (***Figure 6***), and correlated the mean RT within each decile with the corresponding mean PeakLat or PreRespMean, estimated from the average HFA across trials in that decile. We found a significant correlation between PeakLat and RT in a majority of dAI sites for each task (23/40 sites for MCSE, 23/40 for LEC1), while PreRespMean significantly correlated with RT in a large number of sites (16/40 sites for MCSE and 17/40 for LEC1). Overall, this analysis revealed a significant correspondence between RT and both HFA peak latency and pre-response HFA in approximately half of the dAI sites. A second analysis encompassed all iEEG sites recorded in the dASG and dMSG, to test the global effect of reaction time on PeakLat and PreRespMean in that region and not just in selected sites (***Figure 6***). The analysis was conducted separately for each experimental condition, with a similar grouping of correct trials according to reaction time into ten deciles. After computing PeakLat and PreRespMean for each decile, we evaluated the effect of reaction time (i.e decile) on those two variables using linear regression analyses. For the HARD condition of task MCSE, the results indicated that reaction time decile explained 7.65 % of the variation in PeakLat (F(1,1628) = 134.8, p = 2.2e-16), showing that an increase in reaction time was associated with a greater HFA Peak Latency. A similar analysis for PreRespMean revealed a small but significant effect of reaction time decile, explaining 0.2% of the variance (F(1,1628) = 4.8, p < 0.03), indicating a small increase in pre-response HFA with longer RTs. Overall, when repeating the same analysis for the various conditions of tasks MCSE and LEC1 (summarized in ***Figure 6***), we found that the effect of reaction time on preRespMEAN and PeakLat were general, confirming the visual impression derived from ***Figure 3***.

**Figure 6.**
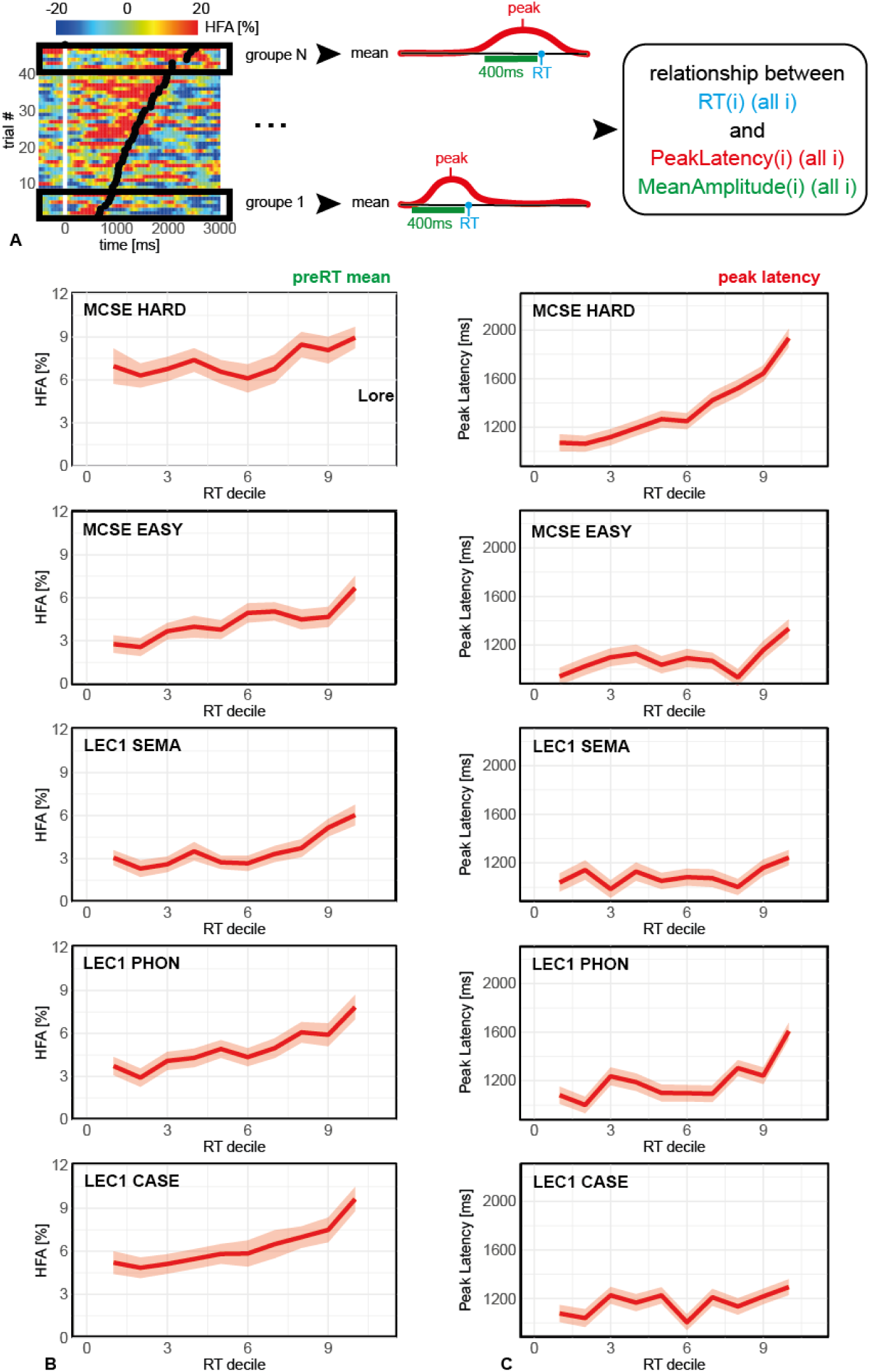
Relationship between dAI response time-course and reaction time. (A) Definition of the metrics used to investigate the relationship between the HFA time-course and reaction time (RT). For a given iEEG site, task and condition, trials were divided into n groups according to reaction time (black rectangles, e.g., reaction time deciles, if n = 10). The average HFA was calculated for each group (red curves) to extract the latency and value of the peak (in red), or the mean HFA in the 400 ms window preceding the reaction time (green horizontal bar), and the mean RT for that group (blue). The analysis investigated the relationship between the mean RT and extracted HFA metrics (peak value, peak latency, pre-response mean HFA), from the cumulative distribution of those measures across all dAI sites. (B) Distribution of the pre-response mean HFA (left, “preRT mean”) and HFA peak latency (right) across all dAI sites, for each reaction time decile (1 = fastest responses, 10 = slowest) and all experimental conditions of MCSE and LEC1. Data were pre-processed as described in panel A (with n = 10). Note the later peak latency and higher pre-response mean HFA for slower reaction times (shaded areas represent the s.e.m. across sites).

### Coincident responses in the dAI and cortical regions supporting task execution

The previous analyses indicated that the dAI response is largely sustained throughout the task independently of the cognitive processes engaged. This finding appears to contradict earlier claims that this region reacts early to incoming stimuli to trigger activity changes in task-negative and task-positive networks. Indeed, Figures ***Figure 7*** and ***Figure 8*** provide clear examples where dAI activations were either simultaneous or later than the responses in those networks. For instance, the dAI response shown for one representative patient during the MCSE-HARD condition (***Figure 8***), overlapped almost perfectly with the response in the intra-parietal sulcus - a major node of the dorsal attention network (DAN). Its time course was also similar to that observed in the cingulate cortex and the middle frontal gyrus, both implicated in attentive visual search. Most notably, the peak of neural activity in the Frontal Eye Field - also part of the DAN - clearly preceded the dAI response. In another patient (***Figure 9***), the increase in dAI activity followed, rather than preceded, the suppression measured in two key nodes of the Default Mode Network during visual search: the posterior cingulate cortex/precuneus and the medial PFC. As illustrated in ***Figure 7***, this tem-poral coincidence between the neural responses recorded in the dAI and in most of the regions involved in the search process was repeatedly observed across patients. Altogether, our observa-tions suggest that the dAI does not consistently react before task-off and task-on networks, and is rather simultaneous with their engagement or disengagement during task-performance. To refine those conclusions, we assessed the precise degree of coincidence between trial-level responses in the dAI and in cortical regions modulated by MCSE using a measure designed to detect long-range functional connectivity from correlated HFA amplitude-fluctuations (see the methods section). The measure estimates the deviation of the distribution of correlation coefficients computed for each trial between the HFA responses measured in two sites, from a surrogate distribution (***Figure 10***). We applied this method to cortical sites exhibiting a significantly HFA increase or decrease during MCSE-HARD and found a widespread distribution of sites significantly correlated with the dAI, with an overrepresentation of executive areas, especially within the dorsolateral prefrontal cortex [17 sites in 7 patients] and a region including the anterior cingulate gyrus and the presupplementary motor area [12 sites in 7 patients] (***Figure 11*** and ***Figure 10***). These findings suggest that the dAI operates in concert with regions associated with cognitive control during active processing of in-coming stimuli. Interestingly, the analysis also revealed significant correlations between the dAI and high-level visual areas in the temporal lobe. This suggests that the dAI might interact with sensory areas during the attentive processing of sensory stimuli.

**Figure 7.**
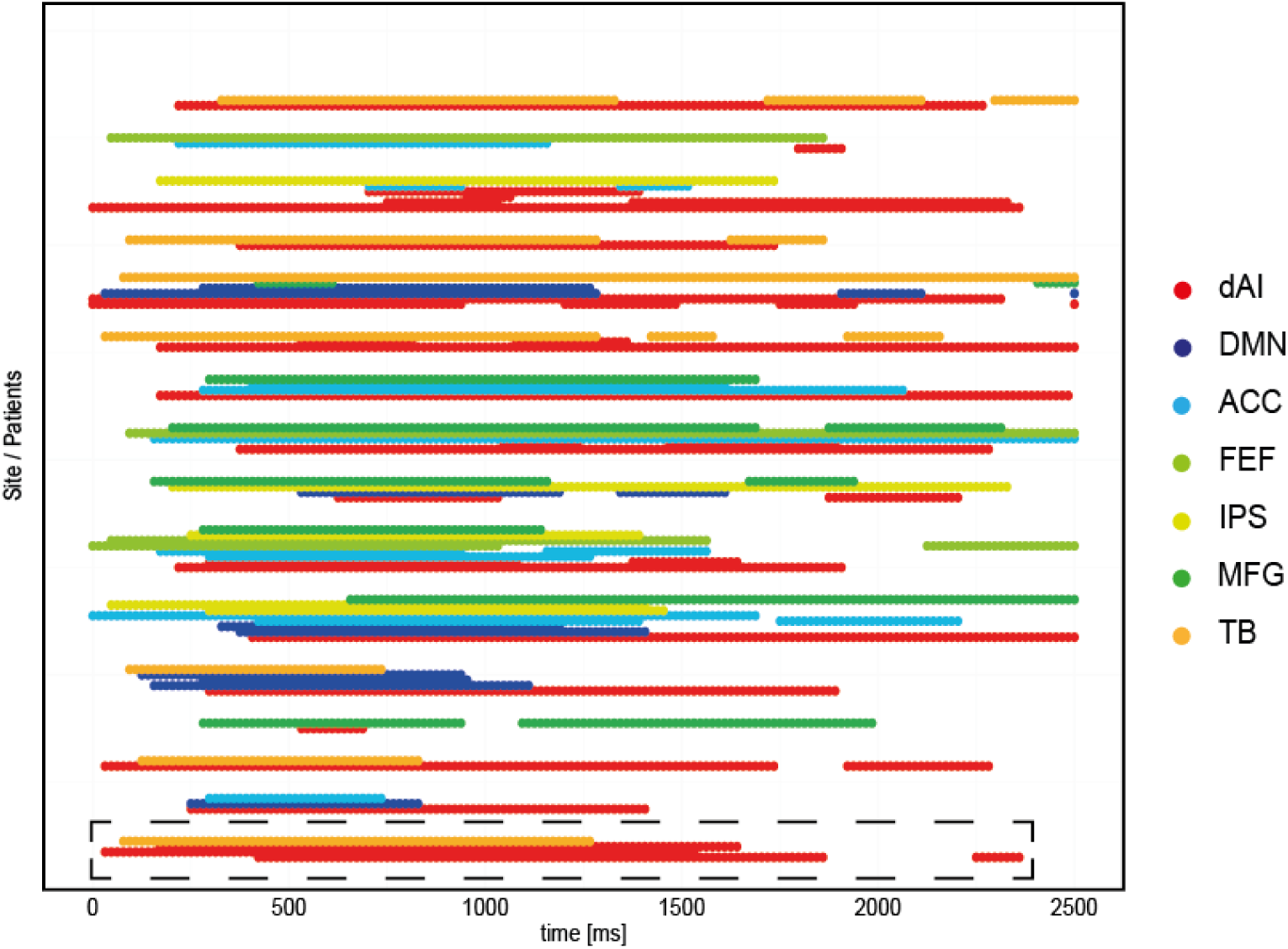
Relative time-course of activation in the dAI and major cortical components of the visual search network across multiple patients. The figure displays periods of statistically significant HFA increase or decrease relative to a pre-stimulus baseline, for selected sites activated or deactivated in the MCSE task (HARD condition) (t-test, p<0.05 uncorrected, discarding isolated periods shorter than 100 ms; see results section for details). Each row represents the time-course of a single site in a single patient, with a vertical grouping of sites belonging to the same patient (i.e. dashed rectangle). Regions displayed include the dorsal Anterior Insula (dAI), the Anterior Cingulate Cortex (ACC), nodes of the Default Mode Network (DMN), the Frontal Eye Field (FEF), the Intra Parietal Sulcus (IPS),the Middle Frontal Gyrus (MFG), and temporo-basal (TB) sites.

**Figure 8.**
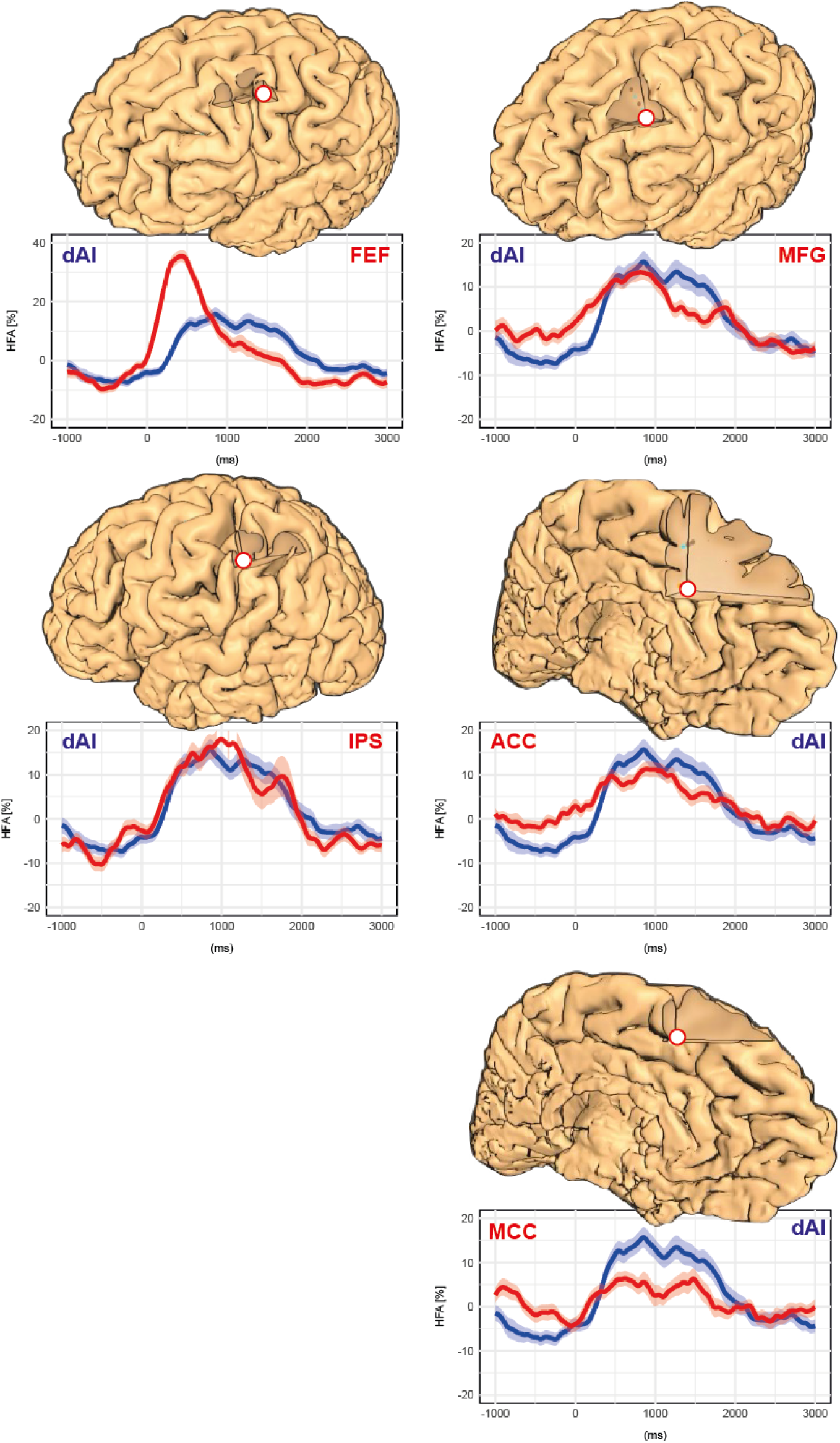
Comparison of the neural response dynamics in key nodes of the visual search network. This figure illustrates the joint response dynamics of a dAI site (in blue) and five cortical sites (in red) active in the MCSE paradigm (HARD condition), recorded in the same patient. Site locations are displayed onto a 3D reconstruction of the patient’s brain and include the Frontal Eye Field (FEF), the Middle Frontal Gyrus (MFG), the Intra Parietal Sulcus (IPS), the Anterior Cingulate Cortex (ACC), and the Mid Cingulate Cortex (MCC). HFA is expressed as percent change relative to the mean HFA measured across the entire recording protocol.

**Figure 9.**
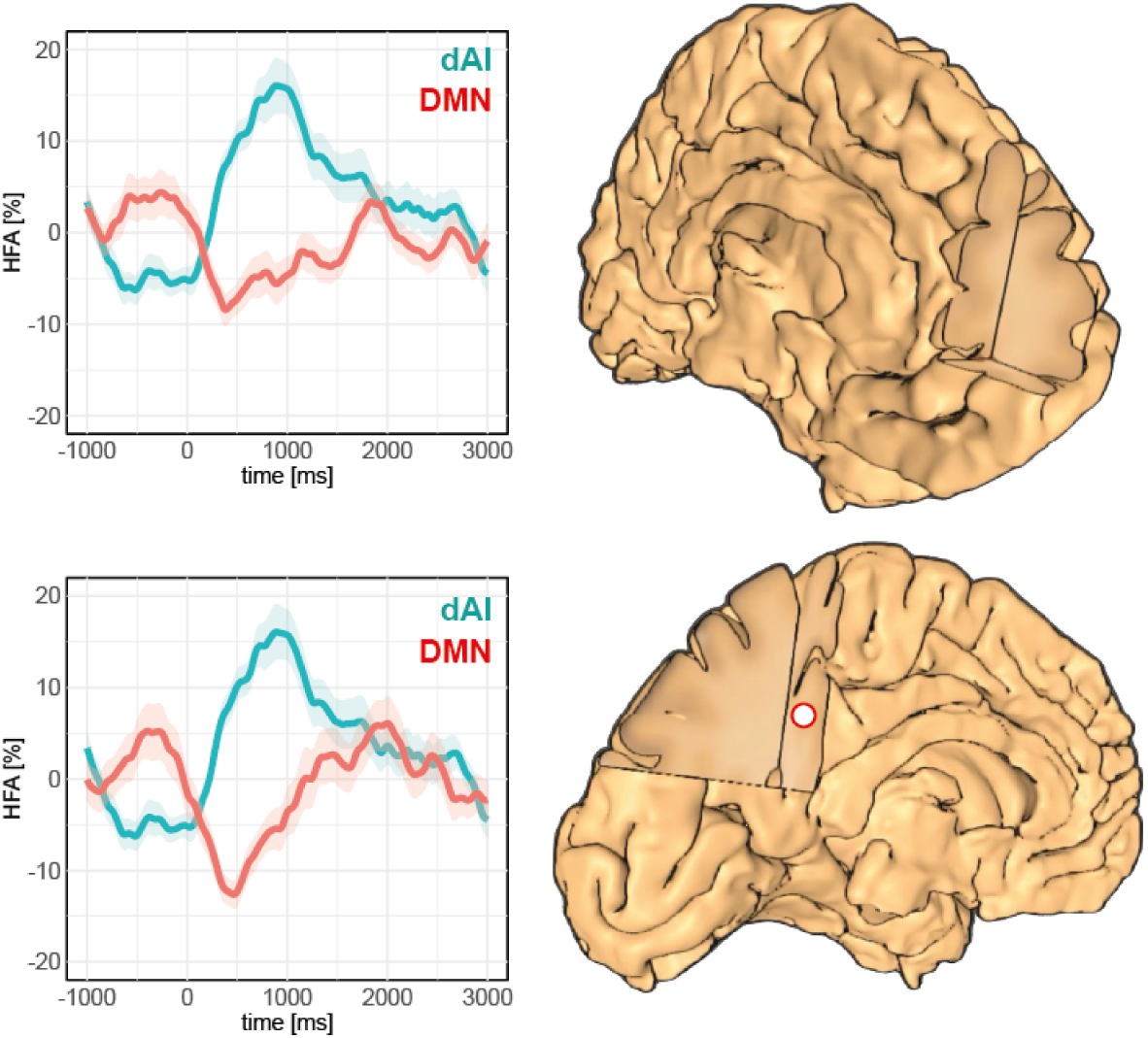
Relative timing of activation and deactivation in the dAI and the Default Mode Network during a visual search. This figure illustrates the temporal dynamics of High-Frequency Activity (HFA, 50-150 Hz) during the HARD condition of MCSE, recorded from a site within the dorsal anterior insular (dAI) (blue curve) and from two nodes of the Default Mode Network (DMN) (top red curve: median Prefrontal Cortex; bottom red curve: posterior cingulate cortex (PCC)/precuneus). Each graph displays the HFA time-course relative to stimulus onset (time = 0 ms) from the same representative patient. Recording Sites are projected upon a 3D reconstruction of the patient’s brain. Note the difference in peak latency for the DMN deactivation and the dAI activation.

**Figure 10.**
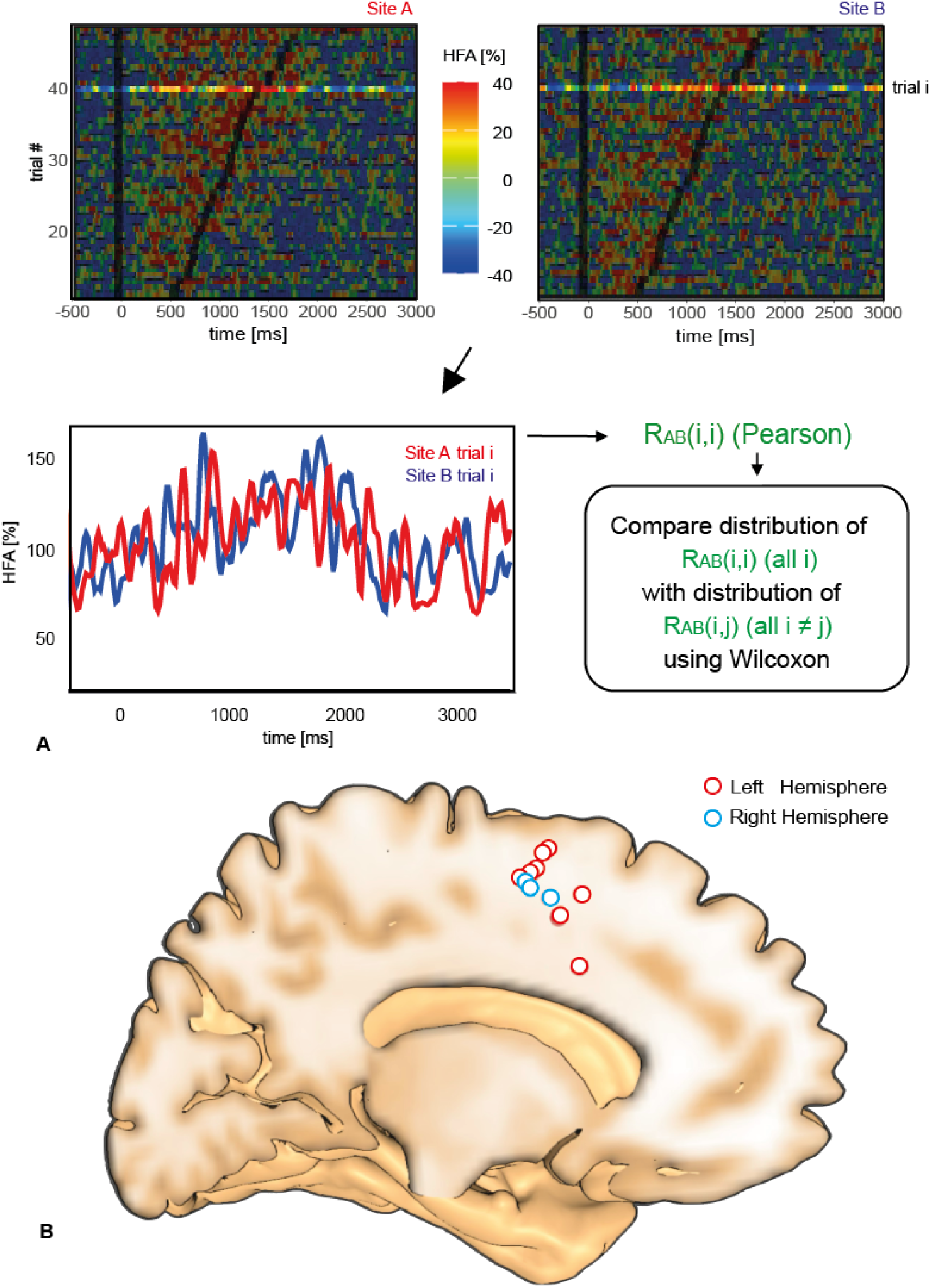
Cortico-cortical functional connectivity assessed by HFA amplitude–amplitude correlations. (A) Illustration of the method used to identify significant correlations between HFA time fluctuations measured in two recording sites (A and B). For each trial 𝑖, the HFA signal is extracted for site A and site B (top matrices), to compute a Pearson correlation coefficient 𝑅_𝐴𝐵_(𝑖, 𝑗). The procedure is repeated for all trials (𝑖 = 1∶nbtrials) to generate the distribution of coefficients obtained when considering the same trial for A and B (test distribution). A surrogate distribution is then obtained by considering all correlation coefficients 𝑅_𝐴𝐵_(𝑖, 𝑗) for all pairs where 𝑖 ≠ 𝑗. The significance of the inter-site amplitude correlation between A and B is then determined by comparing the two distributions with a Wilcoxon test (𝑝 < 0.05, Bonferroni correction). (B) Anatomical location of 11 sites (5 patients) in the cingulate cortex with a significant correlation with the dorsal anterior insula, displayed on a sagittal slice of the Montreal Neurological Institute (MNI) brain template (all sites were projected onto the left hemisphere for clarity).

**Figure 11.**
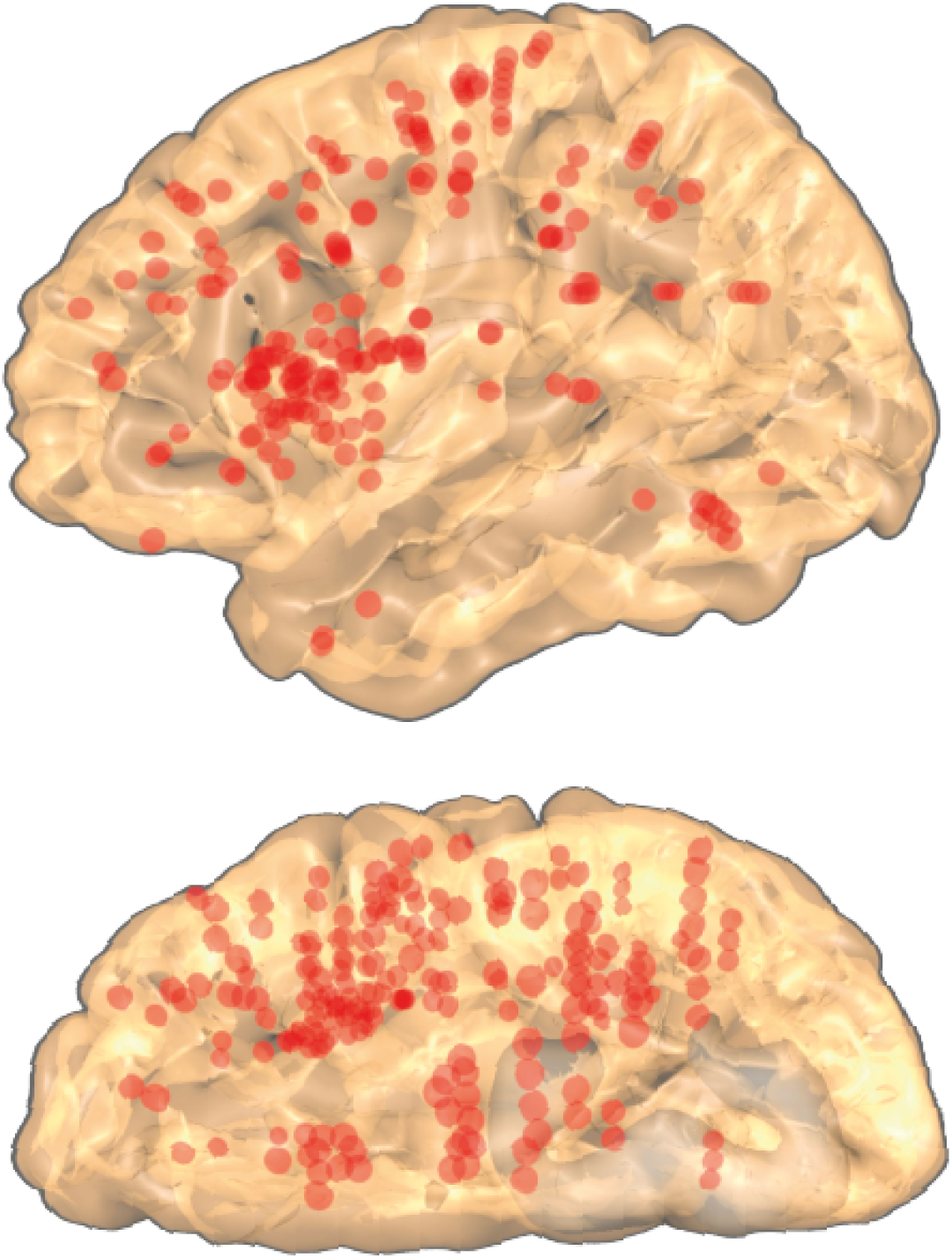
Functional connectivity with the dAI during a visual search task. This figure displays all sites matching two criteria: a) a significant increase or decrease in HFA relative to pre-stimulus baseline during the HARD condition of the MCSE task, b) a significant HFA amplitude correlation with at least one site in the dAI in the same condition (MCSE HARD).

## Discussion

### A shift in viewpoint to characterize the function of the dorsal anterior insula

The objective of the present study was to provide a fine-grained, sub-second characterization of the neural response dynamics in the dAI during a set of cognitive tasks to test implicit predictions of several influential theories about that region. Our methodology was guided by the “shift in viewpoint” proposed by Genon et al. (***Genon et al., 2018***) to characterize the function of a brain region, in which “the a priori defined construct is the brain region and the unknowns are the behavioural functions associated with it”. This strategy shares conceptual similarities with neuroimaging meta-analyses, which aggregate functional data for a specific brain region across multiple cognitive paradigms. However, while typical meta-analyses aggregate data across multiple studies and participant groups—and often conflate activation patterns from neighboring regions (e.g., dAI, vAI, and operculum) due to individual differences in brain anatomy—Genon et al. instead advocated for comparing functional responses across multiple conditions within the same brain (e.g., (***Pinho et al., 2018***)). Our intention was to implement this within-subject characterization using iEEG to record direct neural activity from the same dAI population in individual subjects performing several cognitive paradigms, and to repeat this procedure across a substantial cohort. Importantly, this strategy required careful localization of each iEEG electrode in the native brain space of each patient to avoid the imprecision of automatic labelling (***Duong et al., 2023***).

### A multi-functional signature of the dAI

Several studies have already used iEEG recordings to investigate the functional organization of the insula, particularly in the dAI, and our results are largely consistent with their main findings. Most notably, we replicated the observation by Llorens et al. (***Llorens et al., 2023***) that the dAI responds to task-relevant stimuli with a distinctive time course characteristic of this insular sub-region. Specifically, we observed that high-frequency activity (HFA) between 50 Hz and 150 Hz, in cytoarchitectonic areas Id6 and Id7 (***Quabs et al., 2022***), gradually increased following stimulus presentation and peaked before the behavioral response. Furthermore, the peak latency of the HFA and the amplitude of HFA prior to the response increased with reaction time in the majority of dAI sites. While Llorens et al. (2023) reported this pattern within a single paradigm, our study generalized the finding across highly distinct cognitive paradigms, including an attentional search task and a visual semantic/phonological categorization task.

This illustrates the advantage of recording from the same neural population across multiple cognitive constraints, enabling more comprehensive conclusions than single-task iEEG investigations (***Das and Menon, 2024***). For instance, Llorens et al. (2023) primarily examined the response to probe presentation in a working memory task and concluded that the dAI was involved in decision-making. However, our observation that dAI activity also increased with memory load during the maintenance phase of a similar task suggests a more general role for this region in goal-directed cognition. Additionally, our finding of a robust response to targets in the dAI—approximately 300 ms post-stimulus in a visual oddball paradigm—aligns with and extends previous observations by Citherlet et al. (***Citherlet et al., 2020***) in a comparable task, as well as by Duong et al. (***Duong et al., 2023***) in a visual detection task. But while those authors interpreted this response as evidence of a predominant role of the dAI in detecting infrequent, task-relevant stimuli, the sustained activation during working memory maintenance suggests a broader functional role for this region that transcends mere target detection. Overall, joint analysis of the same neural populations across multiple tasks enables a shift from the classic outside-in approach in neuroimaging (i.e., searching for the neural correlates of a specific psychological function via a tailor-made experimental paradigm) toward an inside-out approach (starting from a brain network and seeking to identify its core functions) (***Genon et al., 2018***).

One result we did not replicate concerns our previous finding that the dAI contributes to error prediction (***Bastin et al. (2017***);(***Billeke et al., 2020***)), as the present paradigms were not designed to test this function. However, the consistent observation of robust dAI responses during successful trials, in the present study, suggests that its role extends beyond error processing. A similar argument applies to recent proposals that the dAI specifically encodes negative valence. Using iEEG, Cecchi et al. reported a functional dissociation between the ventral and dorsal AI in relation to mood and economic decision-making: (a) intracranial electrical stimulation of the dAI reduced patients’ sensitivity to losses and increased risky choices, whereas stimulation of the vAI produced the opposite pattern (***Cecchi et al., 2024***); and (b) iEEG HFA in the dAI and vAI correlated with negative and positive mood, respectively, during a choice task (***Cecchi et al., 2024***). If the allocation of cognitive resources is systematically aversive—except for completely effortless tasks—this might explain why dAI HFA was sustained throughout the stimulus–response interval in all our paradigms. However, an alternative explanation grounded in resource allocation mechanisms explains equally well our findings (see below).

The only other discrepancy with previous iEEG studies concerned Duong’s claim (***Duong et al., 2023***) that Id7 is the only subregion of the dAI that reacts to salient stimuli, as we also observed clear responses to targets in area Id6 during the visual oddball task. However, a careful examination of that study (Figure 5) revealed significant activations within area Id6 as well, warranting further iEEG investigation that would take into account the precise microstructural organization of the dAI. Overall, our findings largely corroborate previous iEEG investigations of the dAI during single tasks, but also provide, we believe, a more comprehensive understanding of this region.

### Conflicting results with previous fMRI findings

In contrast, our findings challenge several conclusions drawn from fMRI studies. For instance, Huang et al. (***Huang et al., 2021***) used a masking and an imagery paradigm to propose that the an-terior insula is “the probable cortical site where conscious access to sensory information is gated,” prior to a global broadcasting of sensory information by the lateral prefrontal cortex (LPFC). Al-though their study broadly situated this gating mechanism in the anterior insula, without distin-guishing between its ventral and dorsal parts, the cortical activation maps shown in their article clearly encompassed the dAI. However, if the dAI were a gate for perceptual awareness, we would expect that (a) stimuli flashed unmasked at the center of the foveal field in our visual oddball task—or in Citherlet et al.’s task (***Citherlet et al., 2020***)—would reliably elicit a response in the dAI (given that participants consciously perceived those images), and (b) that dAI responses to those perceived stimuli would consistently precede those measured in the lateral PFC. Our results, how-ever, appear to contradict both of these predictions. Furthermore, Huang et al.’s proposal is also inconsistent with our observation that, in several dAI recording sites, visual search stimuli failed to elicit a response in participants’ best trials, despite quick and correct performance. One remaining possibility—although not supported by the activation maps shown in their study—is that the vAI, not the dAI, is critical for conscious awareness. In any case, our results should motivate a revision of their hypothesis regarding the role of the anterior insula in perceptual awareness.

Even if the dAI does not determine conscious awareness, most current theories broadly con-verge on the idea that the dAI contributes, in various forms, to a general system involved in de-tecting and reacting to significant external or internal stimuli (such as error predictions) to ensure proper behavioral adjustments (***Centanni et al. (2021***);***Gogolla (2017***);***Molnar-Szakacs and Uddin (2022***)). However, the extent to which the dAI’s involvement spans the entire response process—or is confined to specific stages such as relevant stimulus detection or response monitoring—remains an open question, partly due to the low temporal resolution of fMRI (***Citherlet et al. (2019***);***Citherlet et al. (2020***);***Nelson et al. (2010***);***Power and Petersen (2013***)). In what follows, we show that our iEEG results support a reconciliation of several current functional interpretations of the dAI, provided that some adjustments and extensions are made based on precise timing information.

For instance, a landmark fMRI study is often cited to support the view that the dAI operates only at an early stage, to activate task-positive networks and inhibit task-negative networks (***Sridharan et al., 2008***). While the authors claimed that careful analysis of BOLD signals can reveal subtle latency differences between the dAI and these networks—and thereby demonstrate the primacy and causal influence of dAI activation—our time-resolved iEEG data show no evidence of such dif-ferences. Instead, our findings indicated that deactivation in the default mode network (DMN) and activation in high-level task-positive regions (such as the LPFC) occurred simultaneously with the increase in dAI activity. One might argue that our simple analyses might have failed to capture small latency differences between iEEG signals on the order of a few milliseconds, as well as causal relationships at that timescale; but if such effects were that subtle, they would likely not influence the BOLD signal in a measurable way. Therefore, while the dAI may participate in the switch be-tween task-negative and task-positive networks, it appears to do so continuously throughout the trial, rather than serving as an initial trigger.

The discrepancy between our direct neural recordings and findings from several prominent neuroimaging studies—regarding the relative timing of activation in the dAI and response-related networks—may be attributed to the inherent limitation of fMRI in resolving sub-second timing dif-ferences between consecutive cognitive processes. Once a stimulus is selected, it immediately triggers cognitive processes related to decision-making and response production. This limitation likely contributes to the persistent ambiguity surrounding the role of the dAI—namely, whether it is solely involved in detecting stimuli that require further cognitive processing (i.e., saliency de-tection), or whether it also participates more broadly in subsequent stages, such as initiating or sustaining response-related processes. For instance, an insightful literature review on the dAI concluded that its main function is to act as a gatekeeper to executive control (***Molnar-Szakacs and Uddin, 2022***) —which emphasizes a role in stimulus selection—but also to “prioritize and integrate external sensory information with internal emotional and bodily state signals to mobilize neural resources for optimal executive control,” which implies an active mobilization role throughout task performance.

### Does the dAI belong to the Saliency Network ?

The current consensus identifies the dorsal anterior insula (dAI) as a key node of the Saliency Network (SN), alongside the anterior cingulate cortex (ACC). However, some semantic ambiguity persists regarding the precise definition of “saliency,” and, consequently, the role of that network. For instance, numerous visual search studies equate saliency with a stimulus’s inherent ability to capture attention exogenously due to its unique physical features (e.g., attention capture by a colored singleton among grey items) (***Huang and Yeh, 2011***). In other experimental contexts, however, saliency refers to a stimulus’s contextual relevance—broadly determined by an individual’s current needs, goals, and the risk associated with ignoring that stimulus (***Seeley and others, 2007***). iEEG data obtained during oddball paradigms have clearly shown that the dAI does not necessarily respond to physically salient stimuli (***Citherlet et al., 2019***). Our own recordings revealed that this region failed to respond to rare, high-contrast stimuli presented at the center of the foveal field—stimuli with high feature-related saliency (e.g., pseudowords or faces in a visual oddball paradigm). It follows that the dAI may participate in the detection of salient events, but only in the broad sense used in the original definition of the SN—that is, based on a stimulus’s contextual relevance to the individual at a given moment (***Seeley and others, 2007***). However, our repeated observations that the dAI remains active until task completion across distinct cognitive paradigms call into question its role in the SN. In fact, Seeley (***Seeley, 2019***) suggested that the insular component of the SN is the ventral anterior insula (vAI), whereas the dAI belongs to a separate network originally termed the Cingulo-Opercular Network (CON), involved in the initiation and maintenance of task sets (***Dosenbach et al., 2006***). This view was recently reaffirmed by Dosenbach et al. (***Dosenbach et al., 2025***) in a review redefining the CON as the “Action-Mode Network” (AMN), which is broadly responsible for goal-directed behavior. The authors explicitly differentiated the AMN from the SN and associated the dAI exclusively with the former.

### New insights into the dynamics of the Action-Mode Network

If the dorsal anterior insula (dAI) is a key node of the Action-Mode Network (AMN), our findings provide important insights into the response dynamics of that network during cognitive tasks, as well as its precise functional role. It was originally proposed that activity within the Cingulo-Opercular Network (now AMN) remains stable throughout task performance, maintaining the task set and ensuring that the brain is momentarily specialized for the task at hand through an optimal tuning of functional connectivity (***Dosenbach et al., 2006***). In this framework, the CON/AMN was thought to exert a sustained influence on other regions responsible for processing individual stimuli and generating accurate responses at the trial level—that is, at the timescale of the percep-tion–decision–action cycle. A logical implication is that any interruption of the AMN activity between trials should cause participants to forget the task and make errors. However, the stereotypical response dynamics of the dAI revealed by iEEG—both in our data and in Llorens et al. (***Llorens et al., 2023***)—are characteristic of trial-level rather than task-level activation. Specifically, we observed a progressive increase in activity following stimulus presentation, peaking just before the behavioral response. These results therefore suggest that the dAI contributes to the activation of task-related networks primarily during stimulus processing and response generation, rather than throughout the entire task. If the dAI is indeed a key node of the Action Mode Network, our findings shed a new light on the temporal dynamics—and possibly the functional mechanism—of that network.

### Feeling/filling the need for resources in high-level cortical areas ?

In summary, our multi-task, time-resolved neural recordings support a functional understanding of the dorsal anterior insula (dAI) that encompasses but extends beyond most previous theories. Our findings show that the dAI remains active until the behavioral response in stimulus–response paradigms, as well as during working memory maintenance in the absence of novel stimuli. The dAI must therefore support the entire process leading to the behavioral response at the trial level—not at the task level—which goes beyond mere stimulus detection. Such observations suggest that the dAI might orchestrate responses to task-relevant stimuli through a continuous influence on task-on and task-off networks, rather than serving solely as an initial switch between these networks (***Sridharan et al., 2008***). Although we mostly agree with the proposal that the dAI “mobilizes neural resources for optimal executive control” (***Molnar-Szakacs and Uddin, 2022***), we would add that such mobilization persists until no further processing is required for an accurate behavioral response, making the dAI more than just a “gatekeeper” of executive control. Each time a task is repeated, the dAI allocates resources to the executive regions necessary for that task and possibly reallocates resources away from regions which might interfere negatively (e.g., the default mode network), through its extensive network of connections with cortical regions involved in high-level cognition. Thus, the dAI would act as a continuous “causal hub outflow” (***Molnar-Szakacs and Uddin, 2022***), within the Action-Mode Network.

Within that framework, our observation that the same neural population in the dorsal anterior insula (dAI) is active across multiple unrelated tasks raises an important question: how would those neurons “know” which regions to activate or deactivate? It seems unlikely that the dAI holds a library of pointers toward all possible task-positive networks and uses those pointers to activate the right network ex nihilo on each trial. A more realistic mechanism would exploit the extensive connectivity of the dAI with high-level cortical areas (***Flynn*** (***1999***);***Namkung et al.*** (***2017***)) to detect early activations and deactivations triggered by incoming stimuli and direct resources necessary for task-relevant responses with minimal delay. In iEEG recordings, this would manifest as correlations between amplitude fluctuations in the dAI and connected regions, consistent with our observations. Therefore, one possibility is that the dAI might not participate directly in the response process but rather energize it by mobilizing resources where and when they are needed, akin to sending “reinforcement troops.” Late dAI activation would cause a delay in resource allocation to target areas and increase reaction times, as our data suggest. But that suggestion raises a second question: why would dAI activation occur later in some trials? To reconcile our view with previous reports that the dAI is sensitive to errors and error predictions, we propose that dAI activation might be triggered by a sudden deviation between the perceived and desired cognitive “state” of the individual, as defined in cognitive maps (***Wikenheiser and Schoenbaum, 2016***). When a participant is not making progress toward an immediate behavioral goal, the dAI would receive a go signal to allocate resources selectively to the current task-positive network, consistent with findings that the dAI is active during error awareness (***Klein et al., 2013***). The error signal itself might be computed in neighboring regions, such as the ventral anterior insula (vAI), which reacts to negative feedback (***Billeke et al., 2020***), or the mid cingulate cortex (MCC), involved in performance monitoring (***Fu et al., 2023***). This might explain the strong correlation observed between the dAI and the MCC.

One related possibility is that the dAI might sense the physiological state of specific cortical regions involved in high-level cognition, mostly, thanks to its extensive connectivity network with those regions —much like the posterior insula senses the current and anticipated physiological state of the body (***Livneh et al., 2020***). The dAI would “feel and fill” regional needs for additional resources in cortical areas critical to human higher cognition, and would have evolved disproportionately in our species to serve that purpose. This proposal is consistent with various findings concerning the dAI, including our observation that in a majority of dAI sites, the neural response was weaker for fast and correct responses; which seems to contradict the notion that the dAI supports optimal task completion. Yet, we predict that this minimal engagement of the dAI would occur when resources available in task-positive areas are sufficient to perform the task efficiently, such as when the target captures the participant’s attention exogenously in the visual search task. Our interpretation also clarifies why activity in the anterior insula is strongly correlated with mental fatigue (***Müller and Apps, 2019***) and why participants trained to self-increase activity in their dAI improve performance in demanding tasks ((***Popovova et al., 2024***). Furthermore, we provide a possible explanation for the intriguing finding that electrical stimulations delivered to this region can occasionally generate a sense of bliss (***Villard et al., 2023***). If the dAI receives information about discrepancies between an individual’s desired and actual cognitive states, it might contribute to the conscious experience of dissatisfaction as this gap widens. Conversely, electrical stimulation might prevent the dAI from “feeling the need” to correct the cognitive state, resulting in a perception that the current situation is perfect, with no impetus for action.

### A possible path to effortless attention ?

In conclusion, the activity of the dAI, measured by HFA, might be the most direct measure of an individual’s engagement in a cognitive task. If this is true, our observations that some tasks can sometimes be performed efficiently with little dAI activation suggest the intriguing possibility that we might exert excessive cognitive effort in certain daily-life situations (***Mac-Auliffe et al., 2021***), which offers new perspectives for reducing mental fatigue (***Bruya, 2010***). Additional multi-task iEEG studies are now needed to test this hypothesis across more diverse cognitive situations, including naturalistic settings, as recommended by Genon et al. (***Genon et al., 2018***).

### Limitations

The techniques and methodological choices used in our study have well-known inherent limitations. Most notably, the brains of epileptic patients can only be considered altered models of the healthy human brain. However, well-established guidelines have been developed to enhance the generalizability of conclusions drawn from iEEG studies, including the study of large cohorts of patients with diverse patterns of pathological brain reorganization, as we did here (***Mercier and others, 2022***). Our study also focused on a single component of iEEG signals—high-frequency activity between 50 Hz and 150 Hz—due to its robust relationship with population-level spiking activity and the BOLD signal (***Lachaux et al., 2012***). Further analyses of the same dataset in different frequency bands may yield additional insights about the dAI, especially concerning its functional connectivity. The connectivity measure we employed (amplitude–amplitude correlation) also has limitations, as it does not provide mechanistic insights into the relationships among cortical areas, nor causality information. While several methods exist to extract such advanced information from electrophysiological signals, we were concerned they might occasionally yield ambiguous and unverifiable results, with little possibility to validate them. We believe that the development of intracranial electrode arrays recording from large numbers of individual neurons will offer better options to reveal unambiguous results about the fine interactions between the dAI and surrounding structures. Finally, we acknowledge that our hypothesis may be difficult to distinguish experimentally from the alternative view that the dAI is specialized in processing aversive stimuli (***Cecchi et al., 2022***), especially if detecting a need for resources—or allocating resources itself—is systematically aversive because of its inherent metabolic cost.

## Materials and Methods

### Participants and Recordings

Intracranial electroencephalography (iEEG) recordings were obtained from 169 neurosurgical patients with intractable epilepsy at the Epilepsy Department of the Grenoble University Hospital and the Lyon Neurological Hospital. All participants were stereotactically implanted with multi-lead EEG depth electrodes. Electrode implantations were performed according to routine clinical protocols, based strictly on clinical considerations. Data from any electrode sites within seizure onset zones or showing significant interictal spiking, were discarded from the analysis. All partici-pants provided written informed consent and the experimental procedures were approved by the Institutional Review Board and the National French Science Ethical Committee (CPP 09-CHUG-12, study 0907). All participants had normal or corrected-to-normal vision. Eleven to fifteen semirigid multilead electrodes were stereotactically implanted in the brains of each patient. Those stereotac-tic EEG electrodes have a diameter of 0.8 mm and, depending on the target structure, include 10 to 15 contact leads, each 2 mm wide and spaced 1.5 mm apart (DIXI Medical). Electrode contacts were identified on the individual stereotactic implantation scheme for each patient and anatomi-cally localized using Talairach and Tournoux’s proportional atlas (***Talairach and Tournoux, 1993***). Computer-assisted matching between a post-implantation CT scan and a pre-implantation 3D MRI (VOXIM R, IVS Solutions), or between a post-implantation MRI and a pre-implantation 3D MRI (us-ing IntrAnat, (***Deman and others, 2018***)), facilitated the direct registration of the electrode contacts onto the patient’s brain anatomy. Visualization of iEEG electrodes onto the MNI brain template and individual brains’ anatomy was done using HiBOP (***Vecchio et al., 2024***). Intracranial electroen-cephalography (iEEG) recordings were acquired using a video-iEEG monitoring system (Micromed), enabling simultaneous data acquisition from 128 depth electrode sites. Signals were bandpass-filtered online from 0.1 to 200 Hz and sampled at either 512 Hz or 1024 Hz. During the initial acqui-sition, recordings were made using a reference electrode positioned in the white matter; however, for all subsequent analyses, signals were re-referenced to their direct neighbors using a bipolar derivation to enhance signal quality and spatial resolution of the iEEG recordings (***Mercier and others (2022***);***Ossandon and others (2012***)).

### Spectral analysis

High-Frequency Activity (HFA), between 50 Hz and 150 Hz, was used as a proxy of population-level spiking activity (***Lachaux et al., 2012***). It was extracted from continuous bipolar iEEG signals follow-ing our usual procedure (***Perrone-Bertolotti et al., 2014***): a) signals were first bandpass-filtered into sequential non-overlapping 10-Hz-wide frequency bands (e.g., from 50–60 Hz to 140–150 Hz) using a fourth-order, two-way zero-phase lag Butterworth filter; b) the amplitude envelope for each filtered band was then computed using the Hilbert transform (***Quyen et al., 2001***) and down-sampled to 64 Hz; c) subsequently, the signal was normalized by dividing it by its mean amplitude across the entire recording session and multiplying the resulting signal by 100 (to express activity as a percentage of the overall mean) and d) finally, the normalized amplitude time series from all frequency bands were averaged to produce a single HFA time series, which, by definition, has a mean value of 100 across the entire recording session.

### Experimental tasks and stimuli

#### Visual Search Task (MCSE)

The task was adapted from the classic paradigm of Treisman and Gelade (***Treisman and Gelade, 1980***). It required participants to locate a target character within an array of distractors. Each stimulus consisted of a 6×6 array (36 characters) containing 35’L’ distrac-tors and a single’T’ target, which were randomly arranged. Participants were instructed to locate the target as quickly as possible. To record responses and assess accuracy, participants indicated the vertical location of the target (upper or lower half of the display) by pressing one of two but-tons on a gamepad with their right index finger (for the upper button) or right middle finger (for the lower button). The task included two experimental conditions designed to manipulate search difficulty: in the EASY condition, the target was a gray’T’ presented among black’L’ distractors; in the HARD condition, all characters (both target and distractors) were gray (***Figure 1***). A blocked design was employed, consisting of 8 blocks of 12 stimuli, with 6 stimuli from each condition within each block, presented in pseudo-random order. Each stimulus was displayed for 2500 ms, followed by a 1000 ms interstimulus interval. Stimuli were shown centrally on a 19-inch computer screen posi-tioned 60 cm away from the participant.

#### Language task (LEC1)

The task included three conditions, emphasizing visual, phonological, or semantic processing respectively. In the CASE condition, participants were shown five-or six-letter consonant strings that were incongruent with French graphotactic rules (e.g., “xwxqn”) and were required to indicate with a left index or middle finger press whether the string was written in uppercase or lowercase characters (yes/no response using two buttons). In the PHON condition, participants were asked to judge whether five or six-letter pronounceable pseudo-words contained one syllable or two (with a response input similar to CASE). In the SEMA condition, they were instructed to determine whether five or six-letter words represented living entities (same response apparatus). Participants were instructed to remain silent throughout the task. The task utilized a blocked design, alternating 12 periods, each containing three series of five stimuli from the same condition (CASE, PHON, and SEMA), with the order varied pseudo-randomly between periods. All stimuli were presented foveally on a 19-inch computer screen as black letters on a white background (“Courier” font, 35 mm size), with a black central crosshair stimulus between stimuli. Each stimulus was presented for 2000 ms, followed by an interstimulus interval of 1500 ms.

#### Visuo-spatial Working Memory Task (MVIS)

MVIS is a delayed matched-to-sample visuo-spatial working memory task that tests the retention of spatial dot patterns of varying complexity. Each trial began with the presentation of a central fixation cross (1500 ms), followed by a 4-by-4 grid displayed for 1500 ms, containing two, four, or six dots corresponding to three levels of memory load (LOAD2, LOAD4, and LOAD6) (***Figure 1***). After a 3000 ms retention interval (empty grid), a probe stimulus appeared as a single dot in one of the 16 grid positions. Participants responded by pressing one of two buttons on a gamepad with their right index finger to indicate whether the probe dot’s position was present in the initial stimulus array. The task included 48 test trials (16 trials each for 2-dot, 4-dot, and 6-dot stimuli) and an additional 16 control trials, where a single dot was presented in a different color to indicate that the position did not require memorization. All conditions were assigned randomly across trials.

#### Verbal Working Memory Task (MVEB)

A delayed matched-to-sample task, with the same temporal structure as the MVIS task, was employed to engage verbal working memory. Participants were required to encode and retain the identity of a list of letters over a 3000 ms retention period. The initial list (sample) consisted of a horizontal, central, linear array containing two, four, or six letters, representing different levels of memory load (LOAD2, LOAD4, and LOAD6). Following the retention interval, a single letter was presented as the probe stimulus. Participants responded by indicating whether this probe letter was present in the initial list. The number of trials for each memory load condition was identical to that of the MVIS task design (control trials consisted of the presentation of a single letter in the initial list). Responses were also made via button press with the right index finger on a gamepad.

#### Visual oddball task (VISU)

In this task, participants were instructed to press a button with their right index finger each time a picture of a fruit appeared on the screen (20 targets in total) (***Vidal et al., 2010***). Non-target stimuli consisted of pictures from eight categories: houses, faces, animals, landscapes, objects, pseudowords, consonant strings, and scrambled images (50 different images in each category). All stimuli had the same average luminance, except for pseudowords and consonant strings, which consisted of white letter strings on a black background. All categories were presented within an oval aperture (***Figure 1***) and were presented for 200 ms every 1000–1200 ms in series of 5 pictures interleaved by 3-s pause periods during which patients could freely blink. Visual categories were pseudo-randomised across trials.

### Quantification and statistical analysis

#### Clustering analysis

To prepare for the clustering analysis (***Figure 2***), high-frequency activity (HFA) was averaged for each iEEG site and each trial of the MCSE-HARD condition within five consecutive and non-overlapping windows relative to stimulus onset (W_1_ = [−400 ms ∶ 0 ms] to W_5_ = [1200 ms ∶ 1600 ms]), yielding a 𝑁 × 5 matrix “M” for that site (N is the number of trials). We then: a) applied the Wilcoxon test to each column of matrix M to compare HFA in each time window to baseline HFA (W_1_), b) averaged M across trials to generate a 5-dimensional vector V, c) normalized V to a range of 0 to 100, producing 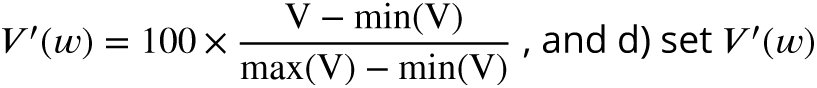 to zero for windows with p-values above 0.001 for the Wilcoxon test (uncorrected). This procedure summarized the HFA dynamics relative to stimulus onset for the MCSE-HARD condition (***Figure 2***). The same steps were repeated for the LEC1-SEMA condition and for both conditions (MCSE-HARD and LEC1-SEMAN) with windows aligned to the motor response RT (W_1_ = [RT − 1000 ms ∶ RT − 700 ms] to W_5_ = [RT + 200 ms ∶ RT + 500 ms], using W_5_ as the baseline). The concatenation of the four 5-dimensional vectors (two for each condition: relative to stimulus onset or to the motor response) produced a 20-dimensional vector for each site, encapsulating the HFA response profile across conditions.

The clustering analysis was conducted on the 20-dimensional vectors of 621 sites using a hierarchical clustering algorithm (complete linkage) in R, specifying three clusters and a distance measure d(V, V^′^) = 1 −R(V, V^′^), where R is the Pearson correlation coefficient. A final visual inspection of the clusters identified potential mislabeling, resulting in cluster adjustments for 4 of the 621 sites.

#### Statistical analysis of High-Frequency Activity (HFA) relative to baseline and across ex-perimental conditions

To detect significant HFA modulations relative to the pre-stimulus baseline, mean HFA within a post-stimulus window of interest (e.g., +500 to 1000 ms relative to stimulus onset) was compared across trials to the mean HFA during a pre-stimulus baseline window (-200 to 0 ms, unless otherwise specified), using a paired Wilcoxon signed-rank test (p<0.05, corrected for multiple comparisons using the False Discovery Rate (FDR) procedure (***Benjamini and Hochberg, 1995***). To compare HFA within the same time window across two experimental conditions (e.g., LOAD6 and LOAD2 for MVIS and MVEB), a non-parametric Wilcoxon test was used (Mann-Whitney U test, p<0.05, FDR correction).

#### Time-stretched grand-average trial matrices

Trial matrixes display HFA as a function of time (x-axis) across multiple trials (y-axis) as a function of reaction time and facilitate the visual identification of stimulus-locked, response-locked, or sustained neural responses (***Ossandon and others, 2012***). Averaging trial matrixes across iEEG sites, within a cortical region of interest, provides a global view of the regional neural dynamics for that region. However, direct averaging is meaningless when iEEG sites have been recorded in multiple patients with different reaction time distributions. To address this, we designed a procedure to normalize the reaction time distribution of each patient to a common, predetermined distribution prior to averaging, and to time-stretch individual trial-matrices at the single-trial level to match that common normalized distribution of reaction times. The procedure proceeded as follows: (a) for each site, the n correct trials were sorted based on their recorded RT, from minimum (RT(1)) to maximum (RT(n)); (b) for each trial i with reaction time RT(i), the HFA time series s_𝑖_(𝑡) for 0 ≤ 𝑡 ≤ RT(𝑖) was stretched to a normalized time series 𝑠*_i_*^′^(𝑡^′^) such that the mapping preserved the activity at stimulus onset (𝑠*_i_*^′^(0) = 𝑠_𝑖_(0)) and at the response time (𝑠^′^(𝑇 (𝑖)) = 𝑠_𝑖_(RT(𝑖))), where 𝑇 (𝑖), 𝑖 = 1,…, 𝑛, was a linearly increasing reaction-time distribution ranging from 𝑇 (1) = 500 ms to 𝑇 (𝑛) = 1500 ms (these values were chosen for clarity). The time-stretching was performed using linear interpola-tion (using the approx function in R): original time points 𝑡 were mapped to normalized time points 𝑡^′^ according to the transformation 𝑡^′^ = 𝑇 (𝑖) × 𝑡 ∕ RT(𝑖). HFA values 𝑠(𝑡) at intermediate time points in the original series were estimated via linear interpolation between measured data points. The resulting time-stretched trial matrices were then averaged across recording sites within each region of interest and subsequently averaged across patients to obtain a grand-average trial matrix for that region.

#### Correlation between HFA and reaction time

For each experimental condition of interest (e.g., MCSE-EASY, MCSE-HARD), correct trials with a reaction time (RT) less than 3000 ms were selected. These trials were sorted by RT and divided into five groups of equal size (corresponding to 20% RT quantiles; see ***Figure 6***. For each group of trials 𝑔 and each iEEG site, three measures were computed: (a) MeanRT(𝑔): the mean reaction time for that group of trials; (b) PeakLat(𝑔): the peak latency of the average high-frequency activity (HFA) across trials within that group; (c) PreRespMean(𝑔): the average across trials of that group of the mean HFA value measured in the 400 ms window that, in each trial, immediately precedes the motor response (RT − 400 ms to RT). For a given set of iEEG sites, the result of this analysis was the proportion of significant correlation coefficients obtained when performing a Spearman correlation between MeanRT(𝑔) and either PeakLat(𝑔) or PreRespMean(𝑔), after pooling all experimental conditions of the task of interest (i.e., 𝑔 = 1∶5× number of conditions). Statistical significance was assessed at 𝑝 < 0.05, uncorrected.

The effect of reaction time on peak latency and pre-response HFA was also assessed at a more global level, for all iEEG sites in the dorsal ASG (Anterior Short Gyrus of the insula) and MSG (Middle Short Gyrus) and for each experimental condition, using linear regression analyses. The procedure ran as follows for a given condition (e.g. MCSE-HARD): a) we selected all correct trials with a reaction time less than 3000 ms; b) we divided the selected trials into ten groups according to reaction time (RT) (ten RT deciles); c) for each decile d, and for every site s of the region of interest, we proceeded as previously described to generate PeakVal(𝑑, 𝑠) and PreRespMean(𝑑, 𝑠) (***Figure 6***), finally, d) we used a linear regression analysis to assess the effect of reaction time (RT decile) on PeakVal and PreRespMean separately (using the ‘lm’ function of R).

#### Functional connectivity analysis

Functional connectivity analysis was conducted using the method described in (***Vidal et al., 2012***). The procedure for two iEEG sites (A and B) and a given experimental condition (e.g., MCSE-HARD) was as follows: for each trial 𝑖, the Pearson correlation coefficient was computed between the HFA measured at sites A and B within a specific time window ([0 ms ∶ 3000 ms] relative to stimulus onset) to produce the distribution of within-trial correlation coefficients 𝑅(𝑖, 𝑖), 𝑖 = 1,…, 𝑛 (where 𝑛 is the number of trials). This distribution was then compared against a null distribution generated by computing the correlation coefficients from different trials for sites A and B, i.e., 𝑅(𝑖, 𝑗) for 𝑖 = 1,…, 𝑛, 𝑗 = 1,…, 𝑛 and 𝑖 ≠ 𝑗. The statistical comparison between the two distributions was performed using an unpaired Wilcoxon test (p < 0.05, after FDR correction across all pairs of iEEG sites). The [0 ms ∶ 3000 ms] window was chosen based on the average reaction time distributions in MCSE and LEC1.

## Acknowledgments

This project has received funding from the European Union’s Horizon 2020 Research and Innovation Programme under Grant Agreement No. 720270 (HBP SGA1) and No. 785907 (HBP SGA2), and by EBRAINS 2.0 (HORIZON-INFRA-2022-SERV-B-01).

---

**Supplementary Figure 1.**
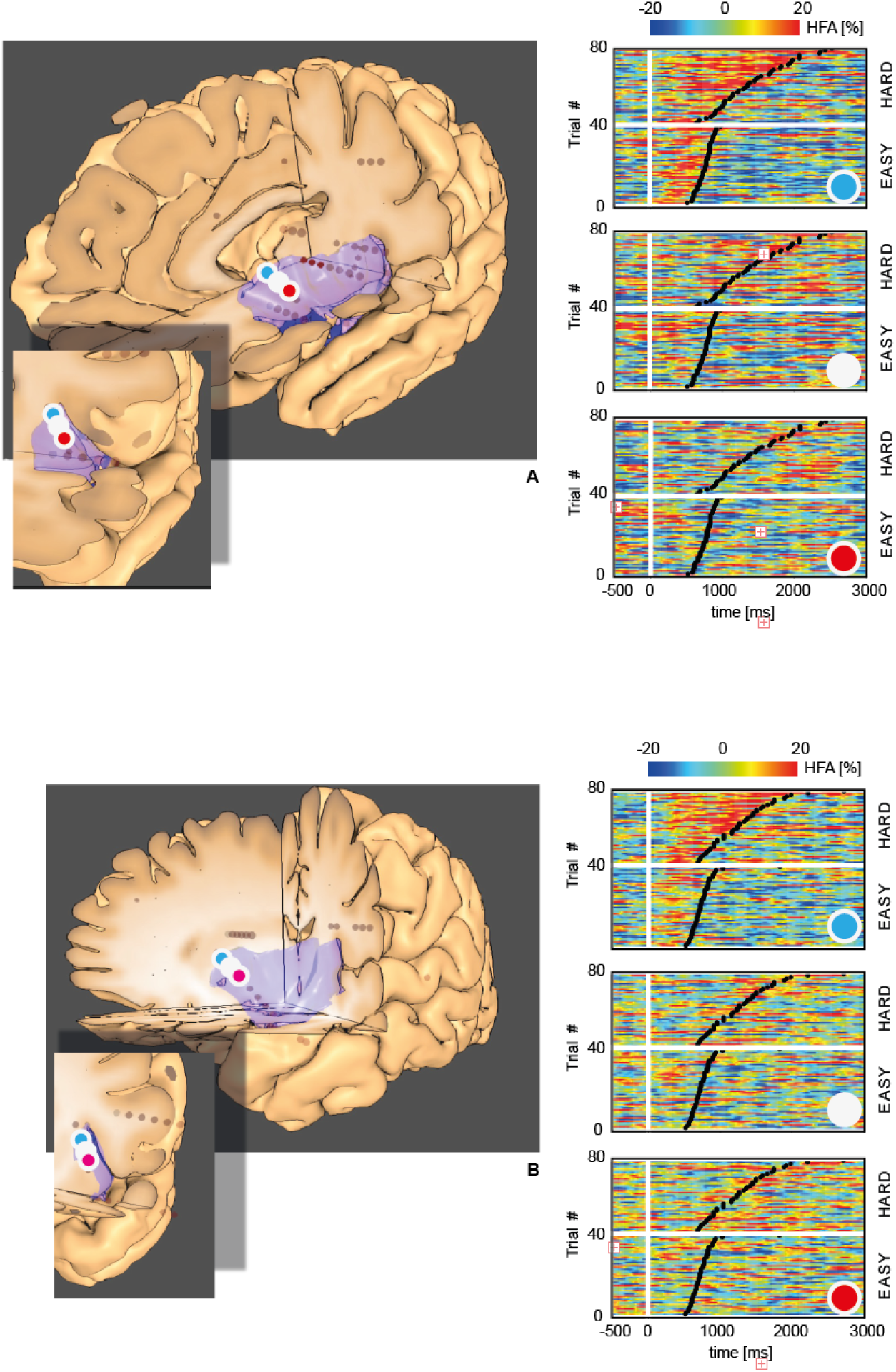
Dorso-ventral functional gradient in the anterior insula during the visual search task. Panels A and B illustrate the variation of the HFA response along a dorso-ventral axis in the anterior insula for two representative patients during the MCSE task, for the two conditions (HARD and EASY). Each matrix depicts the HFA response across latency relative to stimulus onset (x-axis) and individual trials (y-axis), with trials sorted by reaction time (RT), indicated by black dots. In each panel, data are shown for three consecutive recording sites on a single electrode array descending from the superior dorsal anterior insula (dAI) towards the ventral anterior insula (vAI) (inter-site spacing: 3.5 mm). The display suggests a substantial attenuation of the response when descending away from the circular sulcus. HFA is expressed as percent change relative to the mean HFA measured across the entire protocol.

**Supplementary Figure 2.**
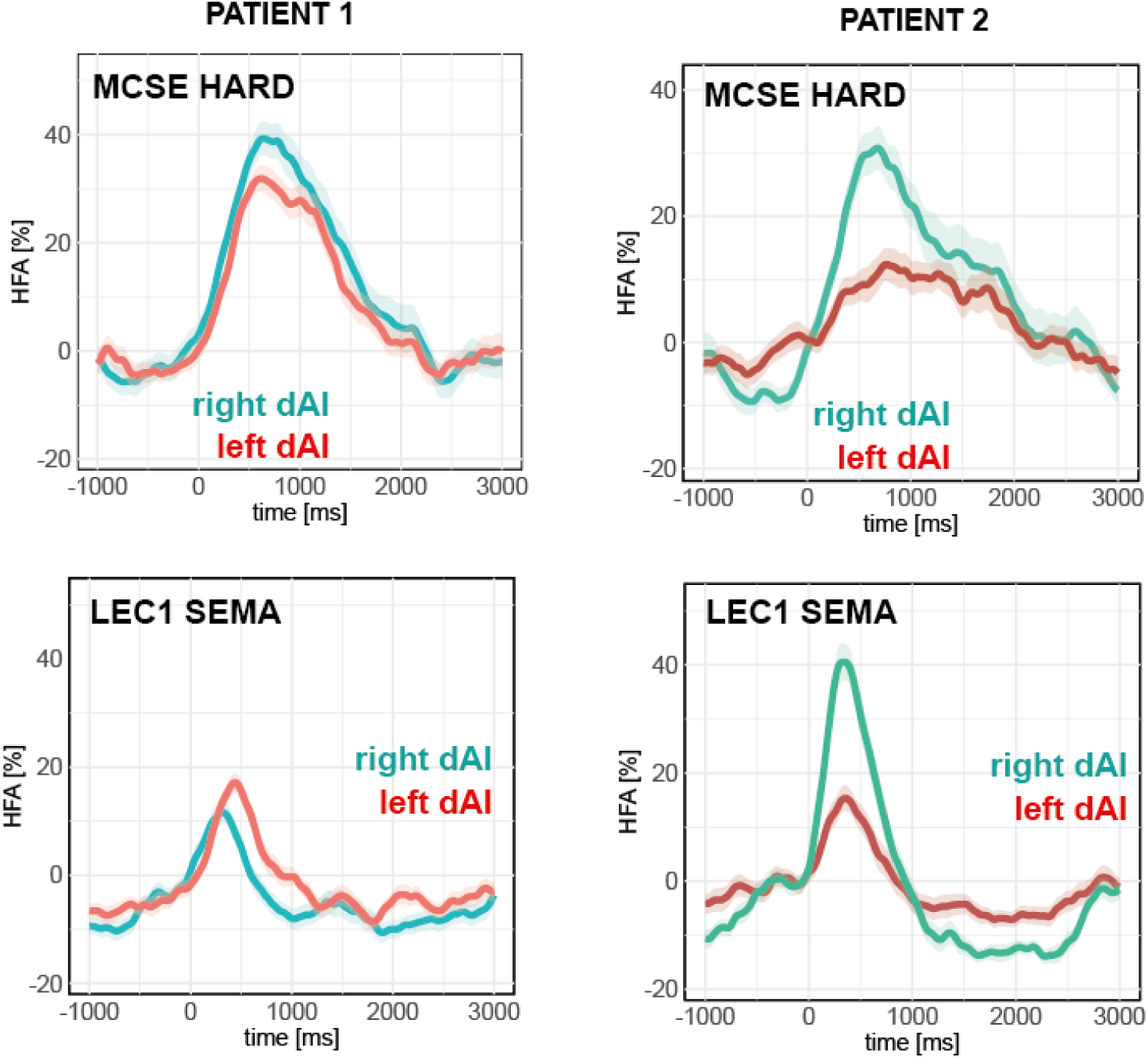
Comparison between response time-courses in the left and right dAI. This figure presents the HFA response in both insulae in two patients with bilateral recordings, for the HARD condition of the MCSE protocol (top row) and in the SEMA condition of the LEC1 protocol (bottom row). Time is relative to stimulus onset and the shaded area indicates the s.e.m. HFA is expressed in % change of the mean HFA measured during the entire protocol.

**Supplementary Figure 3.**
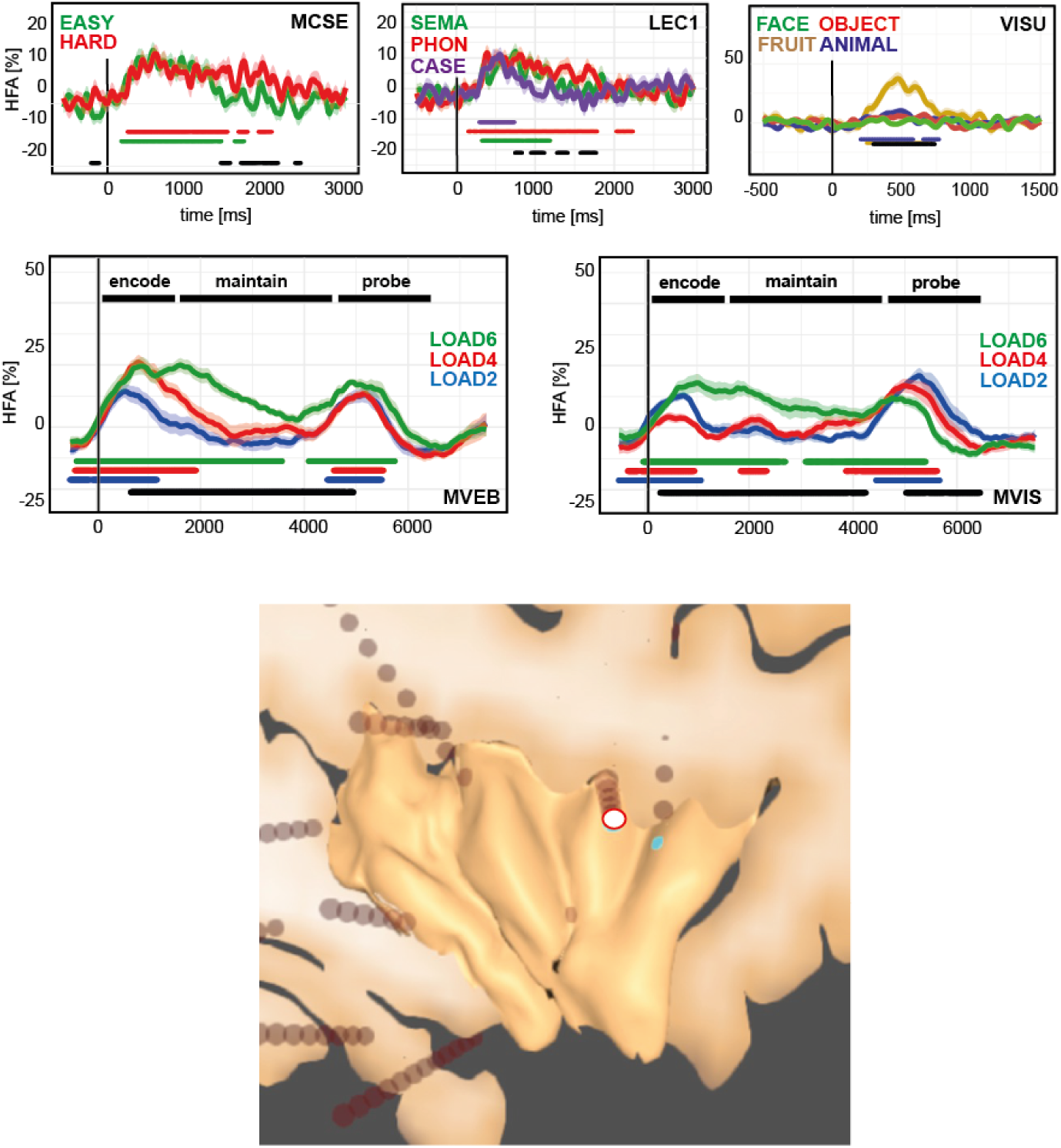
Representative example of the multi-task functional response in the dorsal anterior insula. Similar representation as in ***Figure 5***, for a single site in the superior portion of the short middle gyrus of the insula. Colored horizontal bars indicate time periods during which HFA is statistically higher than baseline (t-test, p<0.05, uncorrected, removing isolated intervals shorter than 100 ms). Black horizontal bars indicate time periods of statistically significant HFA difference between conditions (Kruskal-Wallis test, p<0.05, uncorrected). The white dot indicates the precise location of the recording site onto the participant’s individual anatomy.

